# A non-rectifying potassium channel increases cassava drought tolerance and storage root yield

**DOI:** 10.1101/2025.05.26.655271

**Authors:** W. Zierer, M. Fritzler, T.J. Chiu, R.B. Anjanappa, S.-H. Chang, R. Metzner, J. Quiros, C.E. Lamm, M. Thieme, R. Koller, G. Huber, O. Muller, U. Rascher, U. Sonnewald, H.E. Neuhaus, W. Gruissem, L. Bellin

## Abstract

Cassava (*Manihot esculenta*) is an important crop for food security in the tropics, particularly for smallholder farmers in sub-Saharan Africa, where yields are often severely limited by pathogen pressure, nutrient deficiency, and water scarcity. We expressed a non-rectifying *Arabidopsis thaliana* potassium (K^+^) channel gene version, *AKT2*_var_, in the vascular tissue of cassava plants. The transgenic cassava plants had higher electron transport and CO₂ assimilation rates, a higher bulk flow velocity, and increased source-sink carbohydrate transport, as demonstrated by comparative ^11^C-positron emission tomography and tissue-specific metabolite profiling. Cassava storage root yield was significantly increased in greenhouse experiments and in a multi-year field trial conducted under subtropical conditions. *AKT2*_var_ plants were also more tolerant to drought stress and had higher storage root yield. Targeted alteration of K^+^ transport is therefore a promising strategy to improve cassava productivity without additional fertilizer input and in climate-adverse growing conditions.

## Introduction

Cassava (*Manihot esculenta*) is native to the Amazon region and has become an important staple crop, feeding approximately 800 million people worldwide (*1*). It is particularly important in sub-Saharan Africa (SSA) as a major source of food security and economic support for smallholder farmers. Cassava can grow in poor soils and challenging environmental conditions, but storage root yields are well below its agronomic potential because smallholder farmers in SSA often lack the financial means for fertilizer and irrigation (*1, 2*). Rising temperatures and unpredictable rainfall, leading to more extreme weather events such as droughts or floods, are also negatively affecting cassava production (*3*). Innovative breeding and biotechnology strategies are urgently needed to secure and improve cassava storage root yields in SSA without additional input.

Recent reports have demonstrated a strong relationship between potassium (K^+^) and assimilate (sucrose) transport (e.g. (*4–6*)). K^+^ is an essential macronutrient and a major determinant of crop yield (*7*), and in cassava, storage root yield is a function of crop K^+^ status (*8, 9*). We argue that K^+^ homeostasis can be targeted to increase cassava phloem sucrose transport to storage roots and to improve yield. Phloem transport relies on a solute concentration gradient to generate the necessary pressure to drive mass flow from source to sink tissues (*10, 11*). Although sugars, and particularly sucrose, are the major phloem osmolytes, K^+^ is the most abundant cation in the vasculature and is increasingly recognized as a contributor to phloem mass flow. Phloem K^+^ channels and transporters have been identified (*5, 6, 12–14*) and K^+^ facilitates sucrose loading into the phloem (*15*), most likely by energizing the phloem via a proton-motive force across the companion cell membrane (*5*). The voltage-gated K^+^ channel AKT2 is specific for phloem companion cells (*13*) and functions as an inward-rectifying K^+^ channel (K_in_) to load K^+^ into the companion cell, or, when phosphorylated, as a non-rectifying channel that both loads and releases K^+^ across the plasma membrane (*15, 16*). Circulating K^+^ energy stores, established by phosphorylated AKT2, could increase the efficiency of the H^+^-ATPase-dependent energization of transmembrane phloem loading (*5, 13*). The Arabidopsis AKT2^S210N, S329N^ variant (here AKT2_var_) mirrors the function of the phosphorylated K^+^ channel that, when expressed in Arabidopsis, increases shoot growth, especially under energy limiting conditions (*5*).

Although cassava is characterized by symplasmic phloem loading in leaves (*17*) and symplasmic phloem unloading in storage roots (*18*), the transport phloem still requires active sugar movement to retrieve leaked sucrose during long-distance transport (leakage-retrieval mechanism; (*19*)). Based on the considerable transport distances of cassava and its high K^+^ demand for growth (*8, 9*), we propose that targeting K^+^ transport by vascular tissue-specific expression of AKT2_var_ will facilitate sucrose allocation to growing storage roots, effectively increasing storage root yield (Fig. 1). Here, using non-invasive carbon tracer analysis, we demonstrate that cassava *AKT2_var_* plants have a higher phloem transport velocity and improved agronomic performance in greenhouse and multi-year field experiments, particularly under water-limiting growth conditions.

**Fig. 1.**
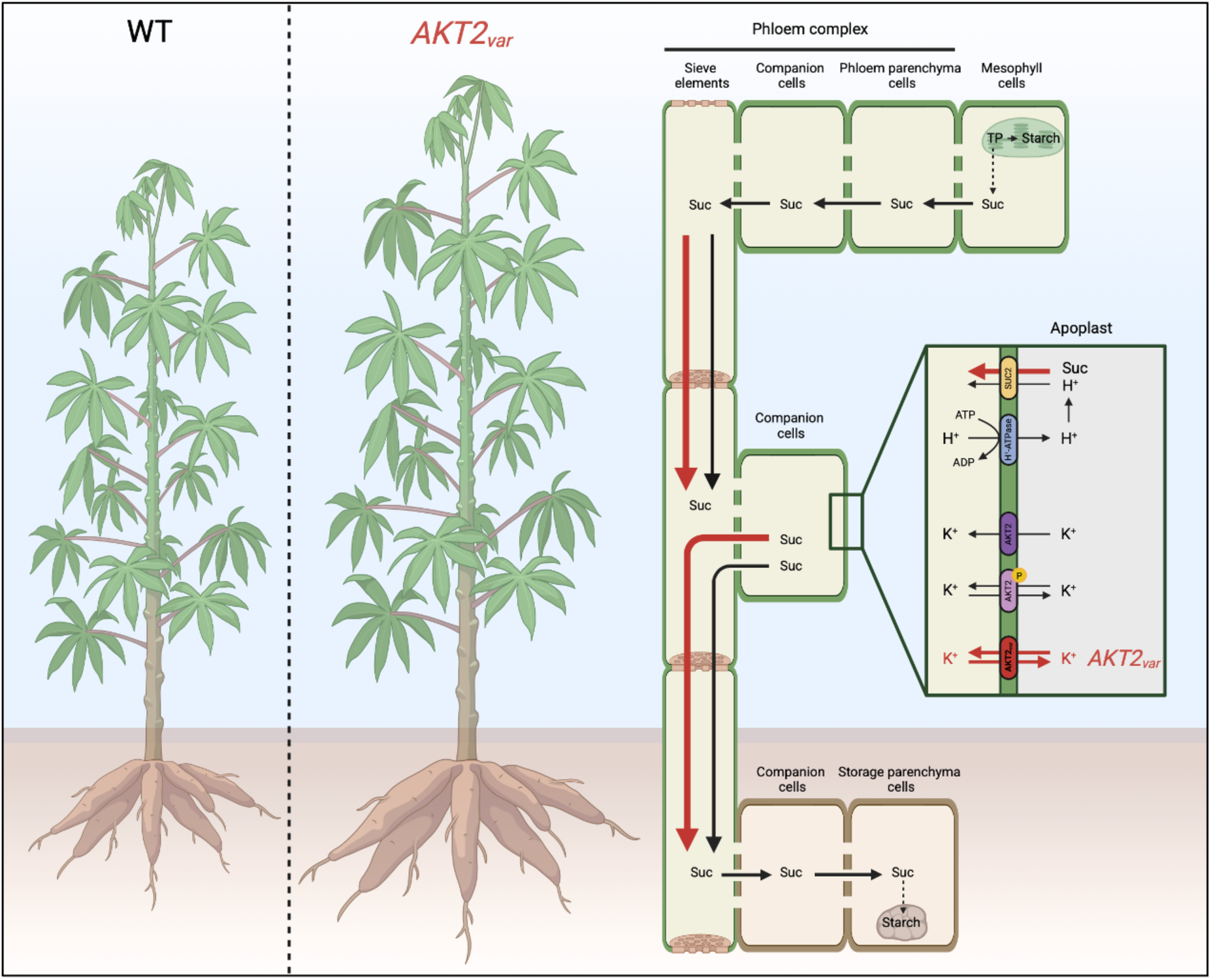
Model of *AKT2_var_* transgene effect in cassava, illustrating its role in increasing phloem transport rates. Black lines and red lines represent K^+^ and sucrose flows in the wild-type and *AKT2_var_* plants, respectively. In addition to endogenous AKT2, either phosphorylated or unphosphorylated, AKT2*_var_* is introduced to modify phloem transport rates in the stem by altering potassium partitioning. This modification is predicted to facilitate sucrose allocation to storage roots by increasing phloem sucrose loading and reloading efficiency.

## Results and Discussion

### The Arabidopsis *AKT2* promoter drives vascular tissue-specific expression of *AKT2_var_* in cassava

We generated independent transgenic lines expressing the *AKT2_var_* gene under the control of the Arabidopsis *AKT2* promoter in the cassava cultivar TMS60444 (Fig. S1A). Four lines had a single T-DNA insertion each (*AKT2_var_*-4255, 4264, 4265, 4266), and two lines had two T-DNA insertions (*AKT2_var_*-4261, 4262) (Fig. S1B). Reverse transcription quantitative PCR (RT-qPCR) analysis using *AKT2_var_*-specific primers revealed a range of mRNA expression levels for *AKT2_var_* in the transgenic lines, with the highest levels in the core tissue of the upper and lower stems (Fig. S1C). Transgenic cassava plants containing the *β-glucuronidase* (*GUS*) reporter gene under the control of the Arabidopsis *AKT2* promoter (*pAtAKT2::GUS*) generally showed GUS expression and a pattern consistent with the mRNA expression data (Fig. S2A). We independently confirmed the RT-qPCR and GUS staining results by end-point RT-PCR using *AKT2*_var_-4266 as an example. As expected, *AKT2_var_* expression was strongest in the different cell types of the stem vasculature, with weaker expression in the vasculature of other plant parts (Fig. S1D). The endogenous *MeAKT2a* and *MeAKT2b* genes are expressed in all cassava tissues analyzed (Fig. S2, B-E), but *MeAKT2a* expression is highest in the phloem tissue, suggesting that MeAKT2a may be functionally equivalent to AtAKT2 (Fig. S1D).

### AKT2_var_ increases phloem transport velocity, facilitating cassava growth and carbohydrate production

Computational modelling had suggested that *AKT2_var_* expression in Arabidopsis promotes the reloading of sucrose leaked from the phloem tissue (*5*), which should lead to higher sucrose concentrations in the sieve elements and companion cells (SE/CC) (*5*) and consequently to higher phloem flow velocity, as proposed nearly 100 years ago (*10*). However, the model prediction has not been experimentally verified in Arabidopsis or tested in crop plants. We therefore used positron emission tomography (PET) to measure phloem flow velocities along the cassava stem in EV (vector control) and *AKT2_var_* lines by incubating leaves with ^11^CO_2_ in the light (Fig. 2A). *AKT2_var_*-4261 and *AKT2_var_*-4262 had significantly higher tracer transport velocities of 11.6 mm min^−1^ (74.5%) and 11.4 mm min^−1^ (70.7%), respectively, compared to 6.7 mm min^−1^ in EV-4234 (Fig. 2B). Photosynthetic fixation capacity was increased in *AKT2_var_* plants with CO₂ assimilation rates of 3.9 µmol m^−2^ s^−1^ (77.3%) in *AKT2_var_*-4261 and 4.4 µmol m^−2^ s^−1^ (100%) in *AKT2_var_*-4262 compared to 2.2 µmol m^−2^ s^−1^ in EV-4234 plants (Fig. 2C).

**Fig. 2.**
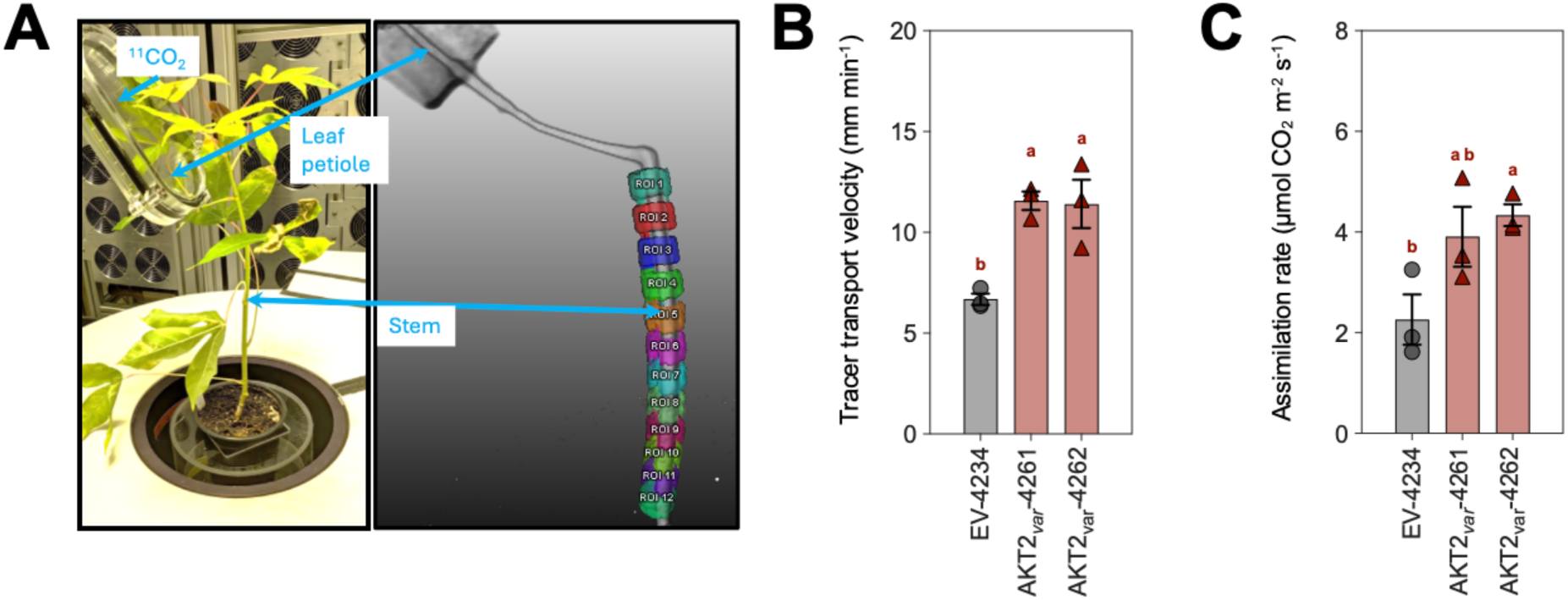
*AKT2_var_* expression increases phloem tracer transport velocities and CO₂ assimilation rates in cassava. (**A**) Example of the experimental setup for the analysis using ¹¹C labelling and positron emission tomography (PET) scanning. The coloured regions of interests (ROI) are used for the velocity analysis. (**B**) Phloem flow velocities in *AKT2_var_*-4261 and *AKT2_var_*-4262 compared to EV-4234 plants. (**C**) Altered CO₂ assimilation rates of ¹¹CO_2_-labelled leaves twelve weeks after planting in the greenhouse. Data in B and C represent the mean of three biological replicates, ± standard deviation. Different lower-case letters indicate statistical significance, as calculated by one-way ANOVA with a post-hoc Tukey HSD test (*p* < 0.05).

The increased transport velocity and assimilation rate in *AKT2_var_* plants had no significant effect on ion (K^+^, Ca^2+^, Mg^2+^, PO_4_^−^, SO_4_^2-^), glucose and fructose concentrations in source (leaf) and sink (stem and root) organs (Fig. S3-4), whereas sucrose and starch concentrations were markedly different (Fig. S4). Leaf sucrose concentrations were reduced by 35% in *AKT2_var_* compared to EV controls, which is consistent with facilitated sucrose reloading and transport in the phloem of *AKT2_var_* plants. A comparable 30% reduction in sucrose levels was also measured in the lower stems of *AKT2_var_* plants compared to EV lower stems, suggesting that the facilitated phloem sucrose reloading capacity provided by AKT2_var_ is not restricted to the leaf phloem tissue. This had a positive effect on starch synthesis and accumulation in *AKT2_var_* storage roots, which had an almost 15% higher starch concentration compared to EV plants (Fig. S4). Starch concentrations were also increased in the lower stems of *AKT2_var_* plants compared to EV controls, which could be potentially beneficial for the typical clonal replanting of cassava from stem segments (stakes) in agricultural production. Taken together, the data suggest that AKT2_var_ can increase long-distance bulk phloem transport of sucrose in cassava while maintaining ion homeostasis, with beneficial effects on carbohydrate accumulation in storage roots and stems. Agronomic evaluation of 19-week-old greenhouse-grown plants of *AKT2_var_* and EV lines showed clear differences in growth performance in four independent cultivation experiments (Fig. 3 and Fig. S5). All *AKT2_var_* plants showed an increase in total dry matter production compared to EV plants. Shoot dry weight (SDW) was increased by 34.0%, 21.2% and 24.6% for *AKT2_var_*-4261, 4262 and 4264 plants, respectively, compared to EV plants (Fig. 3, A and B).

**Fig. 3.**
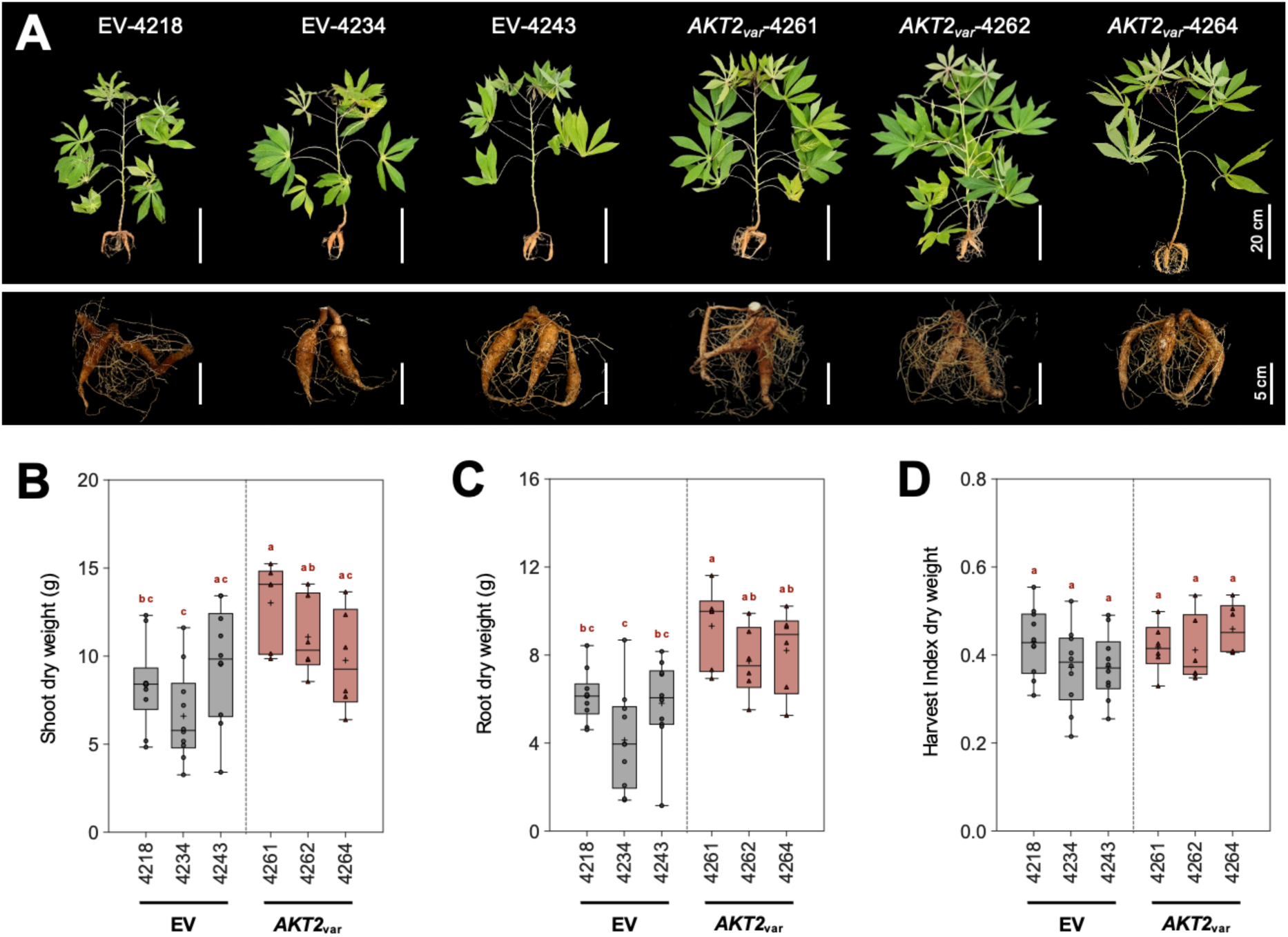
*AKT2_var_* expression results in increased cassava growth rates under controlled greenhouse conditions. (**A**) Representative phenotypes of shoot and root tissues from three vector control (EV-4218, EV-4234, and EV-4243) and three *AKT2_var_* lines (*AKT2_var_*-4261, *AKT2_var_*-4262, and *AKT2_va_*-4264) are shown 19 weeks after planting. (**B**) Shoot dry weight, (**C**) storage root dry weight, and (**D**) harvest index of dry weight. Data in B, C and D represent the mean of 6 to 10 biological replicates, ± standard deviation. In the box plots the central horizontal line represents the median and the plus sign (+) indicates the mean. Different lower-case letters indicate statistical significance, as calculated by one-way ANOVA with a post-hoc Tukey HSD test (*p* < 0.05).

Although cassava storage root development is still at an early stage at 19 weeks, *AKT2_var_* plants already had a more than 55% higher storage root dry weight (RDW) than EV plants (Fig. 3C), which at this growth stage did not affect harvest index (HI) (Fig. 3D).

### AKT2_var_ improves agronomic performance of field-grown cassava

While the data from cassava grown in the greenhouse indicate the positive effect of *AKT2_var_* for plant growth and storage root yield in line with the model assumptions (Fig. 1), they cannot predict the agronomic performance of the *AKT2_var_* plants in the field. We therefore grew plants from the *AKT2_var_*-4255, 4261, 4262, 4265 and 4266 lines together with plants from the EV-4218, 4220, 4221, 4234 and 4243 control lines in three highly replicated randomized field trials (Fig. S6) under subtropical conditions during 2022-2024.

While absolute growth and storage root yield based on agronomic (Fig S7-9) and unmanned aerial vehicle (UAV) measurements (Fig. S10) varied for both AKT2_var_ and EV between experiments, agronomic performance data confirmed a significant positive correlation between shoot and storage root biomass production (Fig. 4). The correlation was less pronounced in 2023 (R = 0.66) than in 2022 or 2024 (R = 0.77), but such variability is not unexpected in the field under varying environmental conditions (Fig. S11). The relative storage root dry matter content (DMC) of all plants in 2022 had a small negative but significant correlation with the fresh weight traits, which was not observed in 2023 or 2024 (Fig. 4A). However, *AKT2_var_* plants still maintained high levels of total storage root dry matter (TRDM) in all three years (Fig. 4B). Spatial correction of the field data using SpATS (*20*) in R and normalization to the mean of EV plant data for each year allowed us to comparatively assess the performance of *AKT2_var_* and EV plants across the independent field experiments and to calculate best linear unbiased estimates (BLUEs) for shoot fresh weight (SFW), root fresh weight (RFW), TRDM, and HI (Fig. 4C). The multi-year statistical analysis confirmed a significant increase in SFW for *AKT2*_var_-4262 and significant increases in RFW and TRDM for *AKT2*_var_-4261 and 4262. HI was also consistently and significantly improved for *AKT2*_var_-4261, 4262 and 4265 compared to EV lines (Fig. 4C). As the growth period during the field trials was eight months compared to the typical 10-12 months for harvestable TSM60444 storage roots and plants were grown from tissue culture plantlets rather than stakes, it is likely that agronomic parameters of *AKT2_var_* plants will continue to outperform EV plants at maturity.

**Fig. 4.**
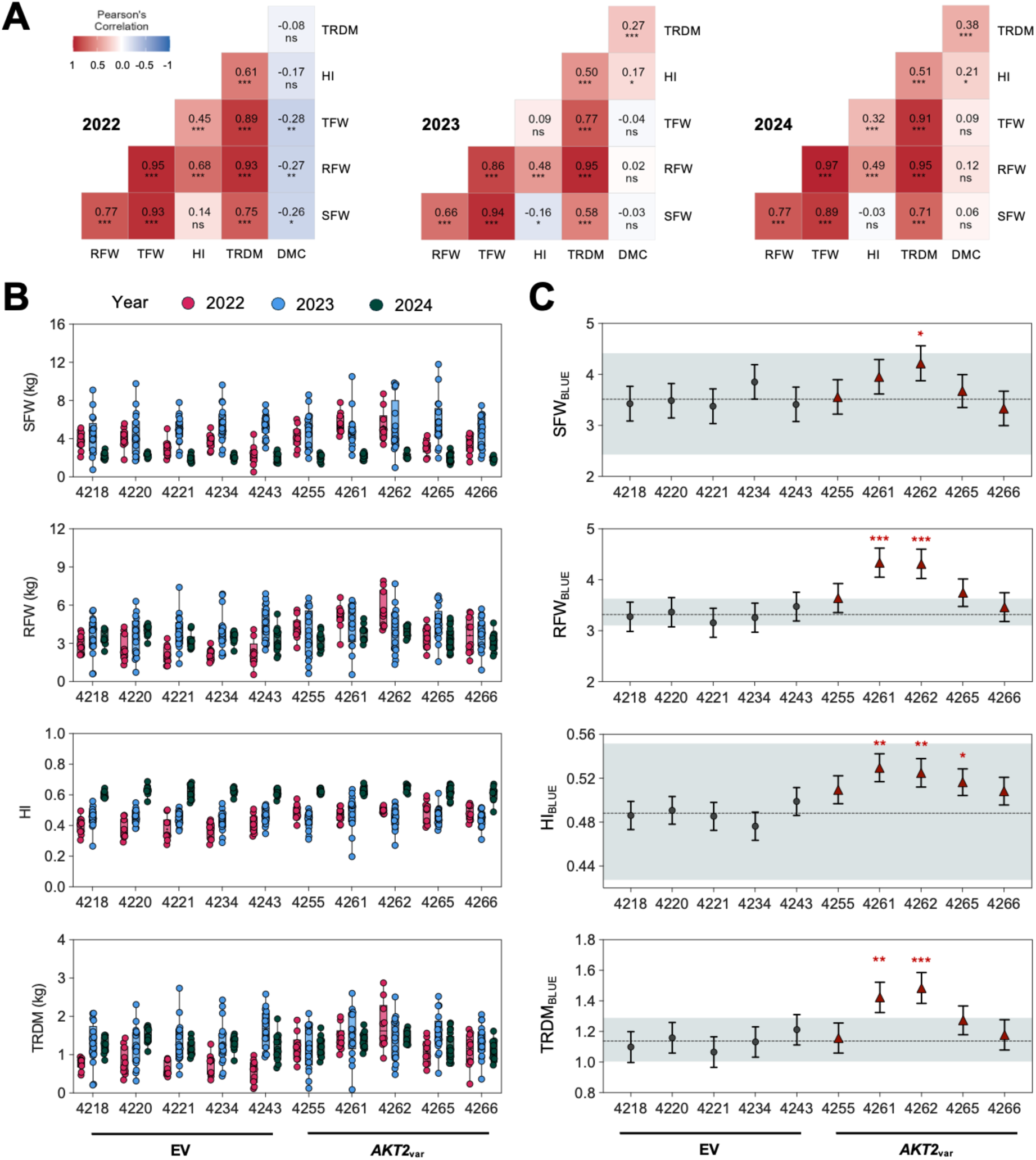
Analysis of multi-year agronomic performance of *AKT2*_var_ and EV control plants in field trials. (**A**) Pearson correlation of agronomic traits per year. Positive and negative correlations are shown in red and blue, respectively. (**B**) Line performance for each year. Spatially corrected data values of replicates (at least ten) are shown for years and traits. (**C**) Line performance after spatial and temporal correction of the raw data (see Materials and Methods). Genotypic best linear unbiased estimates (BLUEs) and standard errors are shown per trait. Significance values are indicated as follows: not significant (p > 0.05, ns), significant (p < 0.05, *), highly significant (p < 0.01, **), and very highly significant (p < 0.001, ***). Traits are abbreviated as follows: SFW = shoot fresh weight, RFW = root fresh weight, HI = harvest index, DMC = dry matter content, and TRDM = total root dry matter.

We next asked whether the improved agronomic performance of *AKT2_var_* field-grown plants was correlated with increased photosynthetic activity and sucrose- and ion partitioning. The electron transport rate (ETR) is a key indicator of photosynthetic performance (i.e., photosynthetic efficiency multiplied by irradiance, saturating at high irradiance as ETR_max_; (*21*)). ETR_max_ was 8% higher in *AKT2_var_* plants compared to EV plants under saturating light conditions in 2022, indicating that AKT2_var_ slightly improves photosynthetic efficiency under field conditions at the early growth stage (Fig. S12). In 2023 and 2024, at a later growth stage, *AKT2_var_* and EV plants showed similar ETR irradiance responses (Fig. S12). *AKT2_var_* field-grown plants had an almost 70% higher K^+^ concentration in the lower part of the stem (Fig. S13), which can explain the almost 40% reduced sucrose concentration (38.5 µmol/g DW) in the lower part of the stem of *AKT2_var_* plants compared to the sucrose concentration (66.9 µmol/g DW) in EV plants (Fig. S14). This difference in K^+^ concentration was not detected in greenhouse plants (Fig. S3), suggesting that AKT2_var_ alters K^+^ homeostasis later in cassava development or under specific environmental conditions. We observed a lower K^+^ concentration in the storage roots of *AKT2_var_*-4261 and 4262 plants, but this difference was not consistent in plants from all *AKT2_var_* lines (Fig. S13).

While other cations (Ca^2+^, Mg^2+^) followed the K^+^ trend in *AKT2_var_* plants, measured anion (Cl^−^, PO_4_^−^, SO_4_^2-^) as well as glucose and fructose concentrations were not significantly different between *AKT2_var_* and EV plants (Fig. S13-14). In addition to the altered K^+^ homeostasis in *AKT2_var_* plants, we found a positive effect on starch accumulation in storage roots, with starch concentrations increased by >50% in the best performing *AKT2_var_* lines (Fig. S14), consistent with the early increase in starch concentrations in the greenhouse plants (Fig. S4). We propose that AKT2_var_-driven K^+^ homeostasis facilitates bulk phloem transport of sucrose from source to sink tissues in the field-grown *AKT2_var_* cassava plants, thereby increasing their starch production and storage root yield.

### *AKT2_var_* plants are resistant to drought stress

The AKT2_var_-dependent increase in phloem flow velocity (Fig. 2C) could also affect the water status of cassava plants, as water availability is closely linked to plant K^+^ homeostasis (e.g., (*22, 23*)). In addition, an increased carbon supply to sink organs could also stimulate the growth of the fibrous root network. We therefore subjected 8-week-old *AKT2_var_* and EV plants grown in the greenhouse to a 5-week drought treatment to assess their growth performance relative to control plants immediately after the drought treatment (intermediate harvest, IH, at week 13) or after a further 5 weeks of normal watering (final harvest, FH; Fig. S15). Under continuous irrigation, *AKT2_var_* and EV plants had similar root system morphology at IH and FH (Fig. 5 and Fig. S16). As previously observed (Fig. 3, 4), all plants from *AKT2_var_*-4261, 4262 and 4264 lines produced almost 30% higher root biomass compared to EV plants (Fig. 4, E and F). Importantly, at IH after drought stress and at FH after rewatering, *AKT2_var_* plants had an increased storage root DW that was >100% higher at FH compared to EV plants, indicating that *AKT2_var_* plants had initiated and maintained their storage root growth during the drought period (Fig. 5, E and F). Stem DW was also increased in drought-treated *AKT2_var_* plants compared to EV plants (Fig. S16). Together this suggests that AKT2_var_ has an especially beneficial physiological effect on cassava growth performance during drought. The levels of the amino acids serine and proline can be used as a proxy for the drought status of plants (e.g. (*24*)), whereas levels of amino acids such as glutamine are typically not altered by this treatment. *AKT2_var_* and EV plants had no significant differences of serine, proline, and glutamine concentrations in leaves and fibrous roots under well-watered conditions. However, serine and proline concentrations specifically increased in EV plants at IH after the drought period, whereas glutamine concentration remained unchanged (Fig. S17). Although proline and serine concentrations also increased in *AKT2_var_* plants at IH after the drought period, they remained significantly lower when compared to EV plants (Fig. S17), indicating that the physiological changes associated with drought stress were more moderate in *AKT2_var_* plants.

**Fig. 5.**
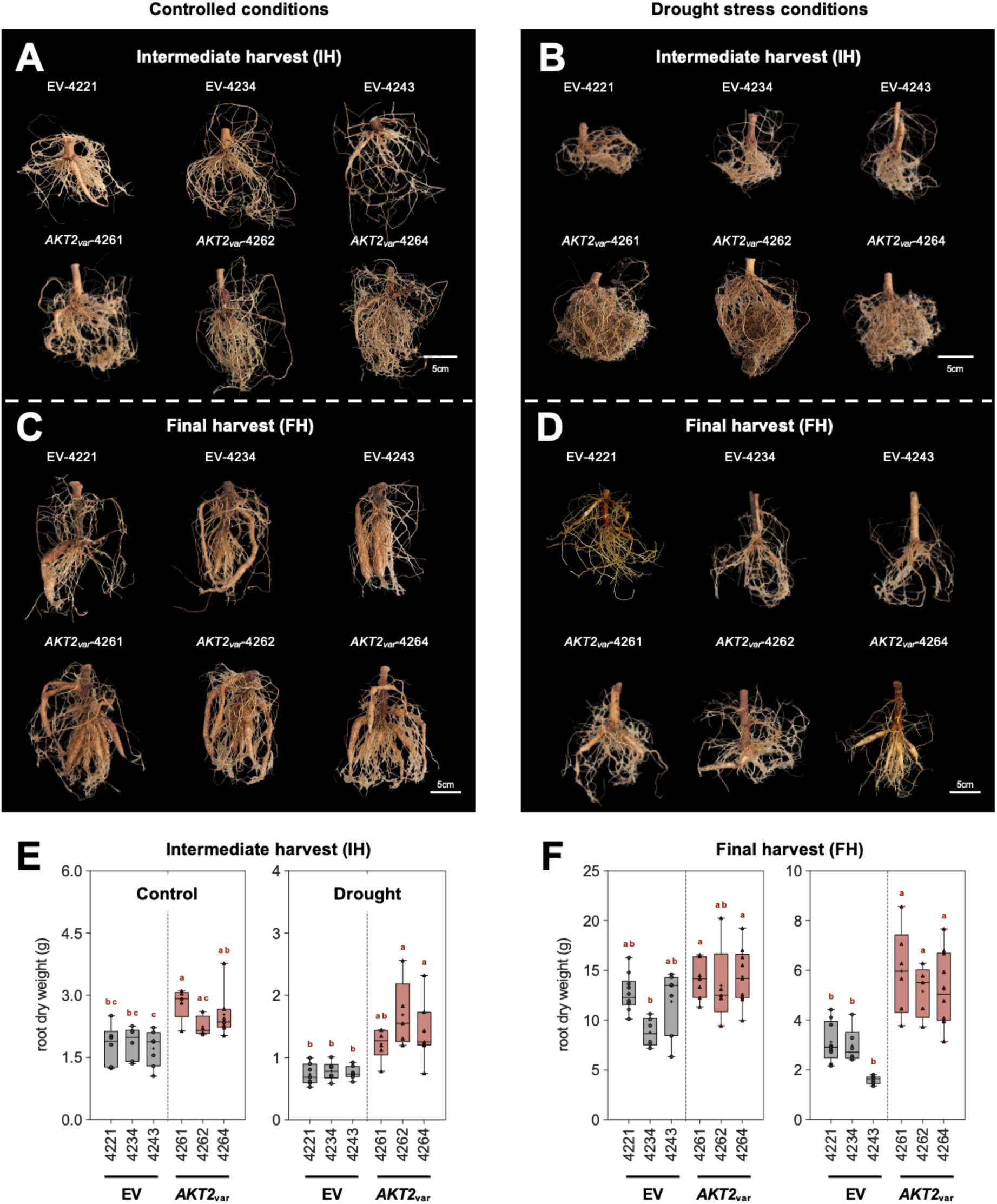
*AKT2_var_* expression significantly improves resistance of cassava to drought stress. To apply drought stress in plants from EV and *AKT2_var_* lines, all plants were first grown under controlled conditions in the greenhouse for eight weeks. This was followed by either a five-week period of drought stress with watering only once a week (drought stress condition), or a five-week period of daily watering (control condition). After these five weeks, an intermediate harvest was carried out for both conditions. Both conditions were then returned to the watering regime of the control condition for five weeks. The final harvest was carried out after this 5-week recovery period. (**A-D**) Representative phenotypes of root tissue from three EV plants (EV-4218, EV-4234, and EV-4243) and three *AKT2_var_* plants (*AKT2_var_*-4261, *AKT2_var_*-4262, and *AKT2_va_*-4264) at intermediate harvest (**A**, **B**) and final harvest (**C**, **D**). Root dry weight was determined at intermediate harvest (**E**) and final harvest (**F**) for both control and drought conditions. Images were digitally extracted for comparison in A-D. In the box plots, the horizontal centre line represents the median and the plus sign (+) indicates the mean. Different lower-case letters indicate statistical significance, as calculated by one-way ANOVA with a post-hoc Tukey HSD test (*p* < 0.05).

The drought treatments of *AKT2_var_* and EV plants had no significant effect on the concentrations of cations (Na^+^, K^+^, Ca^2+^, Mg^2+^) and anions (Cl^−^, NO^3-^, PO_4_^3-^ and SO_4_^2-^) at IH (Fig. S18-19). Sucrose, glucose, and fructose concentrations were significantly increased in *AKT2_var_* plant organs at IH after drought treatment (Fig. S20, B and D), especially sucrose concentrations, which were almost three times higher in stem and root tissues (Fig. S20B). This resulted in up to 50% higher starch concentrations in leaves and stems of *AKT2_var_* plants compared to EV plants (Fig. S20B), suggesting that *AKT2_var_* plants maintained high photosynthetic capacity and metabolism during drought treatment.

## Conclusions

As an essential macronutrient, K^+^ influences the plant water balance by controlling stomatal aperture, thereby affecting photosynthetic efficiency and transpiration rates (*25, 26*), and maintains cell turgor and osmotic balance during abiotic stress conditions (*27*). The results of the present study show that targeting K^+^ homeostasis with AKT2_var_ also increases phloem mass flow velocity, which improves the agronomic performance of cassava under greenhouse and field conditions and even during drought stress. Phloem sucrose transport and K^+^ concentrations are closely linked, as higher phloem K^+^ concentrations can maintain phloem pressure and flow rate when sucrose is reduced in maize (*4*), and phloem K^+^ loading and recirculation between shoot and root affect carbon allocation and grain yield in rice (*6*). Similar to Arabidopsis (*28*), both maize and rice are apoplasmic sucrose loaders (*29, 30*) in which AKT2 orthologs maintain K^+^ homeostasis, and expression of AKT2_var_ channels may also improve their yield, but this has not been experimentally tested. Most importantly, although cassava is a symplasmic sucrose phloem loader (*17*), in which the function of the AKT2 orthologs is not currently understood, our results show that cassava plants expressing the Arabidopsis AKT2_var_ channel have higher phloem transport rate and high photosynthetic efficiency, facilitating source-to-sink transport of sucrose and improving agronomic performance and storage root yield of the crop even during water stress. While the physiological details of the cassava *AKT2_var_* plants require further investigation, the beneficial effects of *AKT2_var_* mutations now provide opportunities to target cassava *AKT2* genes using CRISPR/Cas9 as a breeding tool to improve the crop for smallholder farmers.

## Material and Methods

### Generation of DNA constructs

All binary vectors for genetic transformation were constructed using the Golden Gate cloning toolkit (*31*). The *pAtAKT2::AKT2*_var_ expression cassette contained the following DNA elements: promoter and 5’UTR (1500 bp) of the *Arabidopsis thaliana POTASSIUM TRANSPORTER 2* (*pAtAKT2*) gene, the *Arabidopsis thaliana POTASSIUM TRANSPORTER 2* (*pAtAKT2*) gene coding sequence with S210N and S329N amino acid changes (*AKT2_var_*), a N-terminal 6xHA-tag, and the *Arabidopsis thaliana HEAT SHOCK PROTEIN 18.2* (*HSP18.2*) 3’ UTR. The *pAtAKT2*::*GUS* expression cassette contained the following DNA elements: Promoter and 5’UTR (1500bp) of the *Arabidopsis thaliana POTASSIUM TRANSPORTER 2* (*pAtAKT2*), the modified *BETA*-*GLUCURONIDASE* (“GUSPlus”; (*32*)) gene coding sequence, and the *Agrobacterium tumefaciens NOPALINE SYNTHASE* (*AtuNOS*) terminator. Both plasmids contained the following selectable DNA marker cassette: *Agrobacterium tumefaciens NOPALINE SYNTHASE* (*AtuNOS*) promoter, the *HYGROMYCIN PHOSPHOTRANSFERASE* (*hpt2*) gene coding sequence, and the *CAULIFLOWER MOSAIC VIRUS 35S* terminator. Plasmid sequences are provided in Supplementary Materials.

### Generation of transgenic cassava lines

Genetic modification of the cassava cultivar TMS60444 was carried out as previously described (*33*). *AKT2_var_* and EV cassava lines were generated by transforming friable embryogenic calli (FEC) with *Agrobacterium tumefaciens* containing either the binary vector *p134GG_pAtAKT2::AtAKT2_var_*, which carries the *AKT2_var_* expression and the *hpt2* selectable marker cassettes, or the binary vector *p134GG_Empty vector control,* which contains only the *hpt2* selectable marker. The binary vector *p134GG_pAtAKT2::GUS* was transformed in the same way to generate the *pAtAKT2::GUS* cassava lines. Hygromycin resistant embryos were regenerated for all lines and screened to confirm the presence of the transgene.

### Plant tissue culture

Cassava empty vector (EV) control lines, *AKT2_var_* lines, and *pAtAKT2::GUS* lines were grown in tissue culture containers (Greiner, Frickenhausen, Germany) on MS medium at pH 5.8 (Murashige and Skoog basal salt mixture (MS), including vitamins, Duchefa Biochemie, Harleem, The Netherlands) supplemented with 0.3% (w/v) Gelrite (Duchefa), 2% (v/v) sucrose, and 2 µM CuSO_4_, under sterile conditions. Plants were grown under controlled conditions (16h light/8h dark; 100-120 μmol photons m^−2^ s^−1^, 28 ^°^C in light/26 °C in dark) in a plant growth chamber, before being transferred to soil for greenhouse and field trials.

### Plant culture in greenhouse experiments

Cassava EV control and *AKT2*_var_ transgenic lines were planted either from stem cuttings (stakes) or from sterile culture in standard soil (type ED73, Einheitserde Patzer; Sinntal-Altengronau, Germany) supplemented with 10% (v/v) sand. All plants were grown for 17 weeks in the greenhouse at RPTU Kaiserslautern, Germany (12h light/12h dark; 180 μmol photons m^−2^ s^−1^, 28 ^°^C in light/26 °C in dark). Under standard conditions, plants were watered regularly during the 17-week growth period.

### Drought stress growth conditions

For the drought stress experiments, plants were grown from cuttings and initially grown under standard conditions with regular water supply for the first eight weeks. Periodic drought stress was applied from week nine. Watering was stopped until the soil was noticeably dry, and plants showed phenotypic signs of drought stress, such as leaf wilting and subsequent abscission. After seven days of drought, the plants were watered with a fixed amount of water (100 ml). This watering schedule was repeated for four weeks to simulate periodic drought stress. After the 13^th^ week of growth, the plants were again watered regularly to initiate recovery. The watering schedule is shown in Fig. S15.

### Field trials

Confined field trials were conducted in 2022, 2023, and 2024 at the National Chung Hsing University (NCHU) Experimental Station in Taichung, Taiwan (latitude 24°4’41.50“N; longitude 120°42’56.26“E). For this purpose, plants were imported from sterile culture, transferred to soil, and then hardened in a greenhouse for two months before being transferred to the field. The fields were ridged and covered with black tarpaulin to prevent weed growth. Field trials were conducted in a randomized serpentine design (Fig. S6) with at least 10 replicate plants for each independent transgenic event (line) at final harvest. After a growth period of 8 to 9 months, plants were harvested, and agronomic parameters such as plant height, shoot fresh weight, and root fresh weight were recorded. Dry matter content was determined by drying a representative piece of tissue. Total dry matter content was calculated by multiplying fresh weight by dry matter content. Harvest index was calculated as root biomass divided by total biomass. Samples were collected during plant harvesting and processing as described below. Samples were freeze-dried, processed, and sent to RPTU Kaiserslautern, Germany, for further analysis of ions, sugars and starch.

### Plant harvest and processing of greenhouse-grown plants

All greenhouse-grown plants were harvested after a total growth period of 17 weeks. At harvest, the height of each plant was measured before the plants were separated into three partitions: leaves, stems and storage roots. The weight of each partition was determined, and a tissue sample was collected and immediately frozen in liquid nitrogen for further analysis. When sampling of the stem, part of the stem was separated into peel and core and frozen in liquid nitrogen for further processing. To prevent thawing of the samples, the frozen plant material was ground to a fine powder using a cryo mixer mill (Retsch MM400, Haan, Germany). A sample of 70 mg of the frozen plant material was taken for RNA isolation, and the fresh weight was measured using an analytical scale (Sartorius M-pact AX224, Göttingen, Germany). The plant material was then freeze-dried using a lyophilizer (Alpha 2-4 LDplus, Christ; Osterode am Harz, Germany). After freeze-drying, the dry weight of the plant material was determined using an analytical scale, and samples of 10 mg each were used for analysis of ion, sugar, and starch content.

### Southern Blot analysis

Southern-blot analysis was performed as previously described (*34*) with the following specifications: 10 µg of genomic DNA was isolated (*35*) and separated on a 1% agarose gel. After depurination, denaturation, and neutralization, the DNA was blotted onto a nylon membrane and UV cross-linked. Hybridisation was performed overnight in DIG easy Hyb buffer (Roche, Mannheim, Germany) with digoxigenin-labelled probes (PCR DIG labelling mix, Roche) directed against the *hpt2* coding sequence. After washing and blocking (blocking reagent, Roche), the probes were detected using an alkaline phosphatase-coupled anti-DIG antibody (Roche), CDP-star chemiluminescence substrate (Roche) and a ChemiDoc gel imaging system (Bio-Rad, Feldkirchen, Germany).

### RNA isolation

Frozen plant material from cassava leaves, stem (peel and core) and storage roots were used for RNA extraction. RNA was isolated using the STRN250-1KT Spectrum Plant Total RNA Kit (Sigma-Aldrich, Heidelberg, Germany) according to the manufacturer’s specifications or by using the NucleoSpin RNA Plant Kit (Machery-Nagel, Düren, Germany). The purity and concentration of the extracted RNA was measured using a NanoDrop N60/N50 spectrophotometer (Implen, Munich, Germany) at a wavelength of 260 nm. For subsequent reverse transcription into complementary DNA (cDNA) using the qScript cDNA Synthesis Kit (Quantabio, Beverly, Massachusetts, USA), 600 ng of RNA was used per reaction. Reverse transcription was performed using the Biometra Trio thermocycler (Analytik Jena, Jena, Germany) according to the manufacturer’s specifications. The incubation program used starts with reverse transcriptase (RT) activation at 25 °C for 5 min, followed by reverse transcription at 45 °C for 30 min. The reaction was terminated by heating the samples to 85 °C for 5 min. The resulting cDNA was cooled to 4 °C, diluted 1:6 with H_2_O and stored at −20 °C until use.

### Quantitative RT-PCR analysis

qRT-PCR was performed using the Quantabio SYBR green quantification kit (Quantabio, Beverly, MA, USA) on the PFX96 system (BioRad, Hercules, CA, USA) using specific primers (Table S1). To quantify gene expression, the ΔCq value was calculated by subtracting the Cq value of the gene of interest (GOI) from the Cq value of the housekeeping gene, *MeGAPDH*.

### GUS staining

Various cassava tissues were placed in ice-cold 90% acetone solution. Cross-sections were made manually with a razor blade. These sections were covered with GUS staining buffer (200 mM NaP pH7, 100 mM K_3_[Fe(CN_6_)], 100 mM K_4_[Fe(CN_6_)], 500 mM EDTA, 0.5% SILWET® gold) and thoroughly vacuum infiltrated for 10 min. The GUS staining buffer was removed and replaced with fresh GUS staining solution containing GUS staining buffer with 0.25 mg/ml 5-bromo-4-chloro-3-indolyl-b-D-glucuronic acid (X-Gluc; pre-dissolved in 50 μl DMSO). The GUS staining solution was thoroughly vacuum infiltrated for 10 min. The infiltrated tissues were incubated at 37 °C for 30 min. After removal of the GUS staining solution, 70% ethanol was added to the tissue sections and incubated at 37 °C until the tissues were clear. Images were captured using a Zeiss STEMI SV11 stereomicroscope (Zeiss, Wetzlar, Germany).

### Quantification of ion concentrations

To isolate cations and anions, 800 µl of ddH_2_O was added to 10 mg of lyophilized plant material. The mixture was vortexed thoroughly using a Vortex-Genie 2 (Scientific Industries, Bohemia, New York, USA) and then incubated for 20 min at 95 °C and 500 rpm (Eppendorf Thermomixer Comfort, Hamburg, Germany). After incubation, the samples were again vortexed and placed on ice for 20 min to precipitate the starch. The plant material and starch were then pelleted by centrifugation at 16,000g for 10 min at 4 °C in an Eppendorf centrifuge. 600 µl of the supernatant was transferred to a new reaction tube. Ion samples were diluted 1:5 in ddH_2_O for ion chromatography. Anion and cation concentrations were measured using a 761 Compact IC system (Metrohm, Herisau, Switzerland). A Metrosep A Supp 4-250/4.0 column and a Metrosep A Supp 4/5 Guard/4.0 guard column (Metrohm) were used for the anion measurements. The eluent for anion measurements consisted of 1.8 mM Na_2_CO_3_ and 1.7 mM NaHCO_3_ dissolved in ultrapure water, with 50 mM H_2_SO_4_ as the anti-ion. AMetrosep C4 150/4.0 column and a Metrosep C4 Guard/4.0 guard column (Metrohm) were used for cation concentration measurements, with an eluent of 2 mM HNO_3_ and 1.6 mM dipicolinic acid dissolved in ultrapure water.

### Extraction of sugars and starch

Soluble metabolites, such as sugars, were extracted from 10 mg of lyophilized plant material. Sugars were extracted with 800 μl 80% ethanol. After centrifugation at 16,000g for 5 min, the supernatant was transferred to a new reaction tube, while the remaining pellet containing the plant material was retained for subsequent starch extraction. In preparation for measurement, the supernatant was evaporated using a Speedvac concentrator (Eppendorf, Hamburg, Germany). The resulting pellet was resuspended in 300 μl ddH_2_O. Prior to starch extraction, the pellet from the sugar extraction was washed several times with 80% ethanol and water to remove any residual sugars. After the pellet was washed, 250 µl of ddH_2_O was added, and the samples were autoclaved at 121 °C for 20 min to hydrolyze the starch. For enzymatic starch digestion, 250 µl of a sodium acetate enzyme mastermix (containing 50 U/ml α-amylase, 6.3 U/ml amyloglucosidase, and 200 mM NaOAc at pH 4.8) was added to the pellet and incubated at 37 °C for 4 hours. The digestion was terminated by heating the samples to 95 °C for 10 min.

### Measurement of sugars and starch

Quantification of extracted sugars and hydrolysed starch was performed using an NADP^+^-coupled enzymatic assay as previously described (*36*). For this analysis, the absorbance of NADPH was measured at a wavelength of 340 nm using a photometer (Tecan Infinite M Nano, Männedorf, Switzerland). The sugar concentration was then calculated according to the Lambert-Beer law.

### Measurement of amino acids

For the analysis of free amino acids, 20 µl of an 80% ethanol extract (800 μl) was mixed with 60 µl borate buffer (200 mM, pH 8.8) and 20 µl of aminoquinolyl-N-hydroxysuccinimidyl carbamate (AQC) solution (Synchem UG & Co. KG, Felsberg, Germany; 2 mg ml-1 in acetonitrile). The mixture was immediately vortexed and incubated for 10 min to facilitate derivatization. Quantification of the derivatized AQC amino acids was performed using a Dionex P680 HPLC system with an RF 2000 fluorescence detector (Dionex, Sunnyvale, CA, USA) and a column system consisting of CC8/4 ND 100-5 C18ec and CC 250/4 ND 100-5 C18ec (Macherey-Nagel, Düren, Germany). A gradient of 100 mm sodium acetate/7 mm triethanolamine (pH 5.2, buffer A) and acetonitrile/water (90%, buffer B) (0-100%) was used for amino acid separation. The AQC-derivatized amino acids were detected fluorometrically with an excitation wavelength of 254 nm and an emission wavelength of 395 nm.

### ^11^C-Positron emission tomography (PET) analysis

Two days before and between the PET measurements, the plants were kept in a climate chamber at 28 °C, 65 ± 3% humidity and 400 ± 10 µmol m^−1^ s^−1^ PAR at ambient CO_2_ concentration during the 16h light period. During the 8h dark period, the temperature was lowered to 22±0.5 °C and the humidity was kept constant. The climate chamber containing the PET instrument and the plant during the measurement was set to similar conditions.

On-site production of ^11^CO_2_ was achieved via the ^14^N(p,a)^11^C nuclear reaction by irradiation of N in a gas target with 18 MeV protons at the IBA 18/9 MeV cyclotron of the Institute of Plant Sciences “CYPRES” at the Forschungszentrum Jülich GmbH (Germany). For transfer to the plant labelling circuit, the ^11^CO_2_ was collected in specially designed trapping devices as previously described (*37*). At the end of the collection period, the activity in the closed trap was measured with a collimated scintillation detector (1” NaI Scionix detector, Scionix, Bunik, The Netherlands) connected to an Osprey MCA (Mirion Technologies, Rüsselsheim, Germany) before the trap was transferred to the labelling system. Plant labelling was performed as previously described (*38*). Briefly, the activity in the labelling system was circulated in a closed loop until the target activity of 50 MBq was reached, at which point two valves were switched to include the plant leaf cuvette into the closed circuit for 6 min. After 6 min, the leaf cuvette was switched back to open mode. In open mode, the cuvette was again supplied with conditioned gas from a gas mixing unit, controlled at 26 ± 0.5 °C, 66 ± 4% humidity and 390 ± 10ppm CO_2_, as before the measurement. The outflow from the cuvette was passed through a CO_2_ absorber encased in lead shielding to safely dispose of excess radioactivity. The inflow and outflow of the cuvette was monitored by the following sensors: a differential infrared gas analyser IRGA (LI-7000, LI-COR Biosciences GmbH, Bad Homburg, Germany), a mass flow meter (LowDeltaP, Bronkhorst Deutschland Nord GmbH, Kamen, Germany), an atmospheric pressure sensor (144SC0811BARO, Sensortechnics, First Sensor, Berlin, Germany), and relative humidity and temperature sensor (AC3001, Rotronic Messgeräte GmbH, Ettlingen, Germany). The resulting data were used to calculate the leaf assimilation rate as previously described (*39*). Values are expressed as mean ± standard deviation for a 2-hour period starting 5 min after the end of ^11^CO_2_ labelling. Leaf area was measured destructively after harvest (and used to calculate CO_2_ uptake per leaf area). The gas exchange measurement and labelling system is described elsewhere (*40*). Labelling experiments were performed on the 7^th^ or 8^th^ leaf from the top of the plant, which was the youngest source leaf present on the plant.

The PET system used here, phenoPET, is a custom-built vertical bore instrument for plant measurements with a field of view of 180 mm in diameter and 200 mm in height. Details of the instrument and comparison with other plant-specific PET systems can be found elsewhere (*41*). The system is mounted on a gantry so that it can be moved vertically around a potted plant and the whole setup is installed in a climate chamber. Images were reconstructed from the data using the PRESTO toolkit (*42*).

In the reconstructed 3D images of the tracer signal distribution, cylindrical regions of interest (ROI) were placed along the stem. The position of the ROI in the 3D PET image was determined using anatomical information from visual or imaging observations. The changing activity in these ROIs over time, resulting from ^11^C tracer that was assimilated by the plant after ^11^CO_2_ pulse labelling of a leaf and moving through the stem towards the root, was recorded over time as time-activity curves (*43*). The method used to determine tracer transport velocities from the TAC curves with a compartmental transport model is described elsewhere (*44*). In the part of the stem directly below the petiole insertion (approximately 5-7 cm) connected to the ^11^CO_2_-labelled leaf, the data quality was suboptimal, so that only the lower part of the stem in the field of view was used for transport velocity analysis.

### Photosynthesis measurements

In 2022, photosynthesis data were collected from *AKT2_var_* and EV plants in June at the confined field trial at NCHU Experimental Station, Taichung, Taiwan, using a Mini-pulse amplitude modulation (PAM) system (MiniPAM; Heinz Walz, Germany). Measurements were taken on three top leaves under naturally fluctuating light conditions (Fig. S21). Electron transport rate (ETR) was calculated following (*45*) and fitted to an exponential rise-to-maximum curve to derive ETR_max_ at saturation, indicating photosynthetic capacity (*21*). In 2023 and 2024, photosynthesis data were collected in late November and mid-October, respectively, using a Monitoring-PAM system (MoniPAM; Heinz Walz, Germany; Fig. S21) configured to automatically record measurements every 15 minutes during daylight hours. ETR and ETR_max_ were computed as described above. Instrument examples are shown in Fig. S21, A and B and measured plant locations for each year are shown in Fig. S21, C and E.

### Plant height measurements by unmanned aerial vehicle (UAV)

UAV flight campaigns were carried out using an Okto-XL 6S12 microcopter (HiSystems GmbH, Moormerland, Germany). A high-resolution RGB camera (Sony alpha 6000 with 35 mm lens; Sony Group Corporation, Tokyo, Japan) was used to collect images with 80% overlap (side and front) at 27 m above ground level (mAGL), resulting in a pixel size of 0.003 m. Nadir images were collected near solar noon, typically between 11:00 and 13:00 h local time, on a weekly or bi-weekly basis. A total of 30 ground control points were used to georeference the data based on their known position measured by a real-time kinematic global navigation satellite system system, achieving an accuracy of approximately 0.03 m. Individual raw images were further processed using the photogrammetric structure of the motion software Metashape (Agisoft LLC, St. Petersburg, Russia), from which georectified mosaic images and digital elevation models (DEMs) were generated. From the DEM the crop surface models (CSM) were calculated providing plant height information in meters above the ground level for each UAV data acquisition. Plant height per plant was calculated as the 95^th^ quantile of the CSM values within an approximately 0.50 m buffer around each plant center, aiming to reduce noise from outliers (*46*). Plant volume was estimated as the sum of the CSM pixel values within each buffer, multiplied by the pixel area.

### Statistical analysis of field data

Data were processed in a stepwise manner to account for spatial and temporal variation. First, the data were corrected for field design and spatial trends by trait and year using the R package SpATS (*20*) according to the formula:

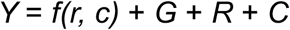

*Y* represents the phenotypic value, *f(r, c)* is a smoothed bivariate surface defined over rows and columns, *G* is the genotype effect, *R* is the row effect, and *C* is the column effect. The number of spline points was set to two thirds of the total number of rows and columns. Based on the spatial correction, outliers were excluded if the residual was more than three standard deviations from the mean.

The best linear unbiased estimates (BLUEs) and residual errors were retained as spatially corrected values (*47, 48*), which were further used for correlation and temporal analyses. In a second step, a linear mixed model was fitted to account for the temporal variation between the years. The previously spatially corrected values were used to calculate the genotypic BLUEs across the years for each trait using the R package lme4 (*49*) and lmerTest (*50*) according to the formula:

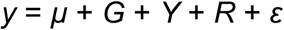

Y represents the spatially corrected value of the respective trait, *µ* is the overall mean, *G* is the fixed effect of the genotypes, *Y* is the random effect of years, *R* is the random effect of the replicates, and *ε* is the residual error. Prior to modelling, an averaged EV control was calculated from the individual EV controls, which served as the reference group in the t-test.

## Acknowledgments

FAU: WZ, CL, and US thank Aaron Engelmann, Birgit Peterson, Eva Düll, Ingrid Schießl, Michaela Reiser, Otilia Ciobotea, Sanika Lakshman for technical support. MT thanks Lukas Kronenberg for support on the field trial analysis.

RPTU: LB, MF, and HEN thank Adrian Heide, Verena Müller, Frank Reinhard, Regina Jerhof, and Maike Siegel for technical support and sample preparation.

NCHU: WG thanks Yong-Shen Gu and Guan-Bo Wang for their field trial support.

FZJ: RK, RM, and GH thank A. Chlubek, E. Breuer, M. Dautzenberg and D. Pflugfelder for technical support with PET. JQ, OM, and UR thank Juliane Bendig and Anna van Doorn for UAV support.

## Author contributions

Conceptualization: HEN, US, WG, WZ

Methodology: CEL, GH, JQ, LB, RBA, RM, SHC, WG, WZ

Investigation: CEL, GH, JQ, LB, MF, MT, OM, RM, SHC, TJC, WZ

Visualization: GH, JQ, LB, MF, MT, RM, TJC, WZ

Funding acquisition: US, WZ

Project administration: US, WZ

Supervision: HEN, LB, OM, RK, UR, US, WG, WZ

Writing – original draft: HEN, LB, WZ

Writing – review & editing: LB, WG, WZ

## Supplementary Materials

### Supplementary Figures

**Fig. S1.**
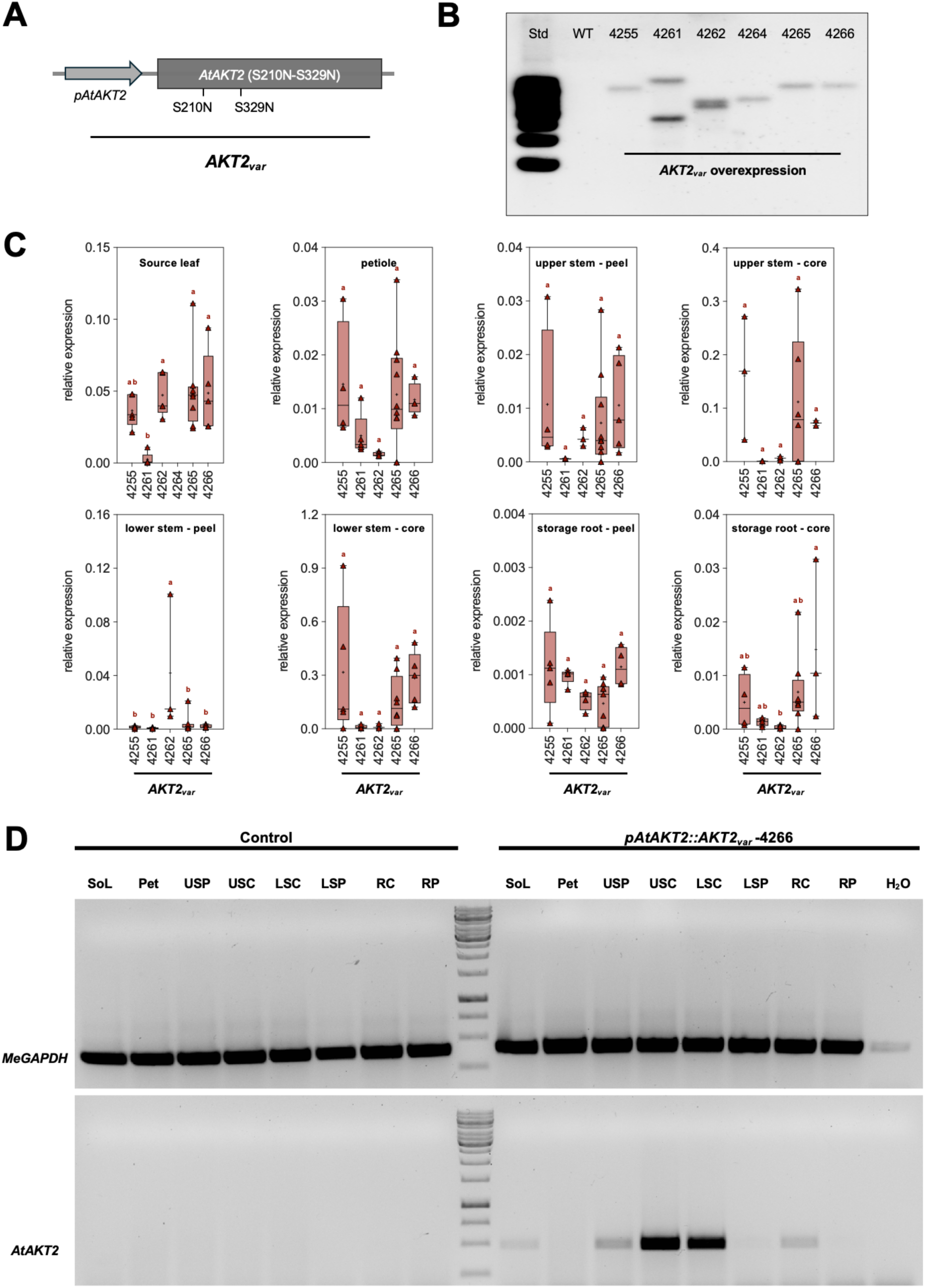
Expression of *AKT2_var_* under control of the Arabidopsis *AKT2* promoter results in expression in different cassava tissues. (**A**) Schematic representation of the mutant *AtAKT2* gene (S210N-S329N, *AKT2_var_*) fused to the Arabidopsis *AKT2* promoter that was transformed into cassava. (**B**) Southern Blot analysis of *AKT2_var_* transgenic lines (*AKT2_var_*-4255, 4261, 4262, 4264, 4265 and 4266) compared to a wild type (WT) plant. (**C**) Quantitative real-time PCR measurements of relative *AKT2_var_* mRNA expression levels in leaf, petiole, tuber, upper, middle, and lower stem, and storage root after normalization to *MeGAPDH*. The horizontal line in the box plots represents the median and the plus sign (+) indicates the mean. Different lower-case letters indicate statistical significance, as calculated by one-way ANOVA with a post-hoc Tukey HSD test (*p* < 0.05). (**D**) Semi-quantitative real-time PCR measurements of relative *AKT2_var_* mRNA expression levels in source leaf tissue (SoL), petiole (Pet), upper stem peel (USP) and core (USC), lower stem peel (LSP) and core (LSC), and root peel (RP) and core (RC) with the corresponding *MeGAPDH* loading control of *AKT2_var_*-4266. The PCR was performed with 35 cycles. Primers “AKT2_CDS_RT_F1/R1” were used (Table S1).

**Fig. S2.**
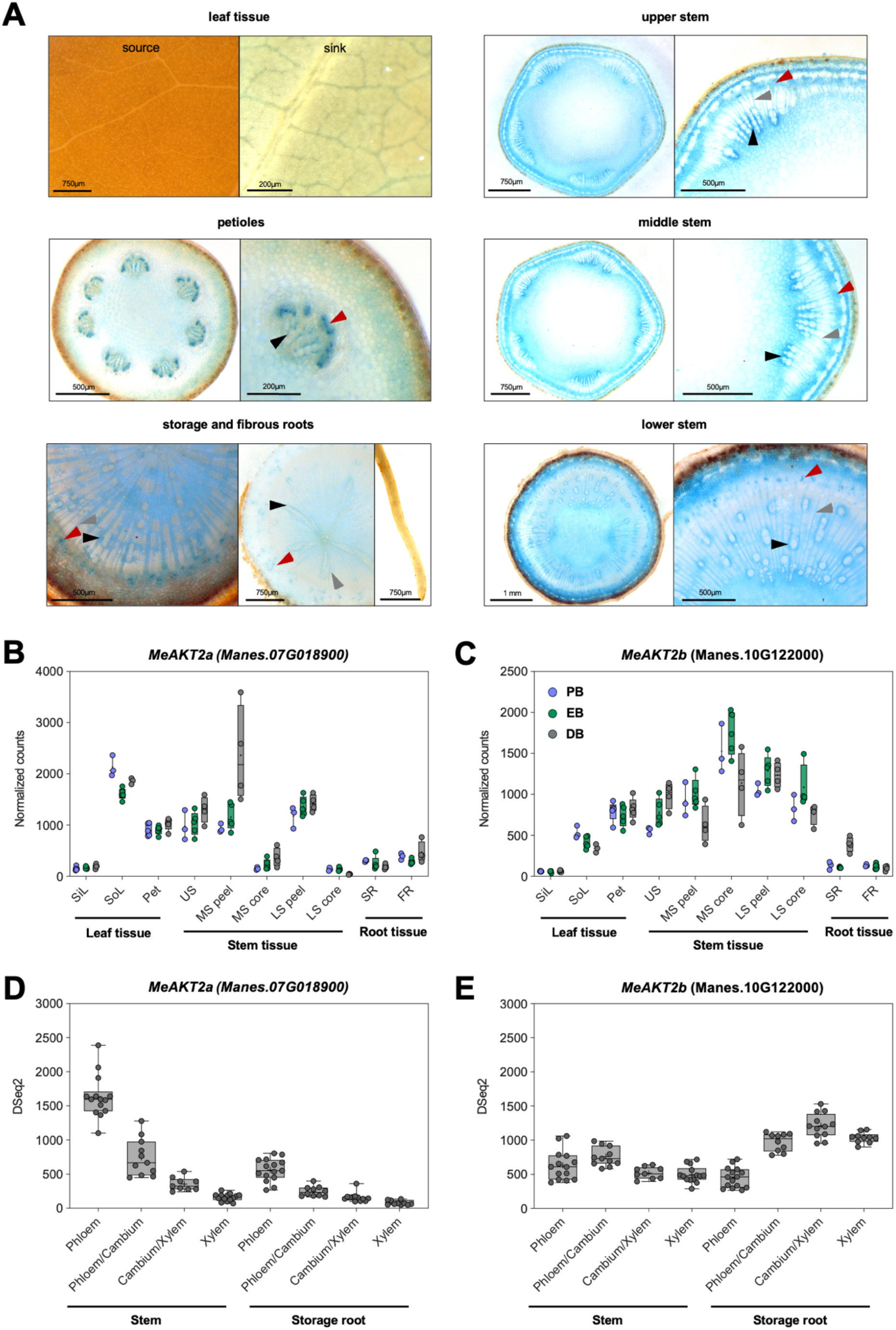
Activity of the Arabidopsis *AKT2* promoter and expression *MeAKT2a* and *MeAKT2b* in cassava. (**A**) Histochemical GUS staining patterns of representative *pAtAKT2::GUS* transgenic cassava lines. Shown are source and sink leaves, cross sections of petioles, upper-, middle-, and lower stems, as well as the storage root and fibrous roots with root tip. Upper, middle, and lower stem are defined as the green part of the stem near the apex (about 5 cm below the apex), the transition zone between the green stem and the brown/grey stem, and the base part of the stem, respectively. Red arrowheads mark phloem companion cells (dotted structures). Black arrowheads mark xylem parenchyma cells closely associated with xylem vessels. Grey arrowheads mark xylem ray cells connecting phloem and xylem. (**B-C**) RNA expression profiles of *MeAKT2a* and *MeAKT2b*. DSeq2-normalized transcript data was published previously (*17*). PB = pre-bulking stage, approximately 30 days after planting (DAP), EB = early bulking stage, approximately 45 DAP, DB = during bulking stage, approximately 60 DAP. SiL = sink leaf, SoL = source leaf, Pet = petiole, US = upper stem, MS peel = middle stem peel, MS core = middle stem core, LS peel = lower stem peel, LS core = lower stem core, SR = storage root, FR = fibrous root. (**D-E**) Expression profiles across the radial dimension of stem and storage root for *MeAKT2a* and *MeAKT2b*. DSeq2-normalized transcript data was published previously (*51*). The horizontal line in the box plots represents the median and plus (+) the mean.

**Fig. S3.**
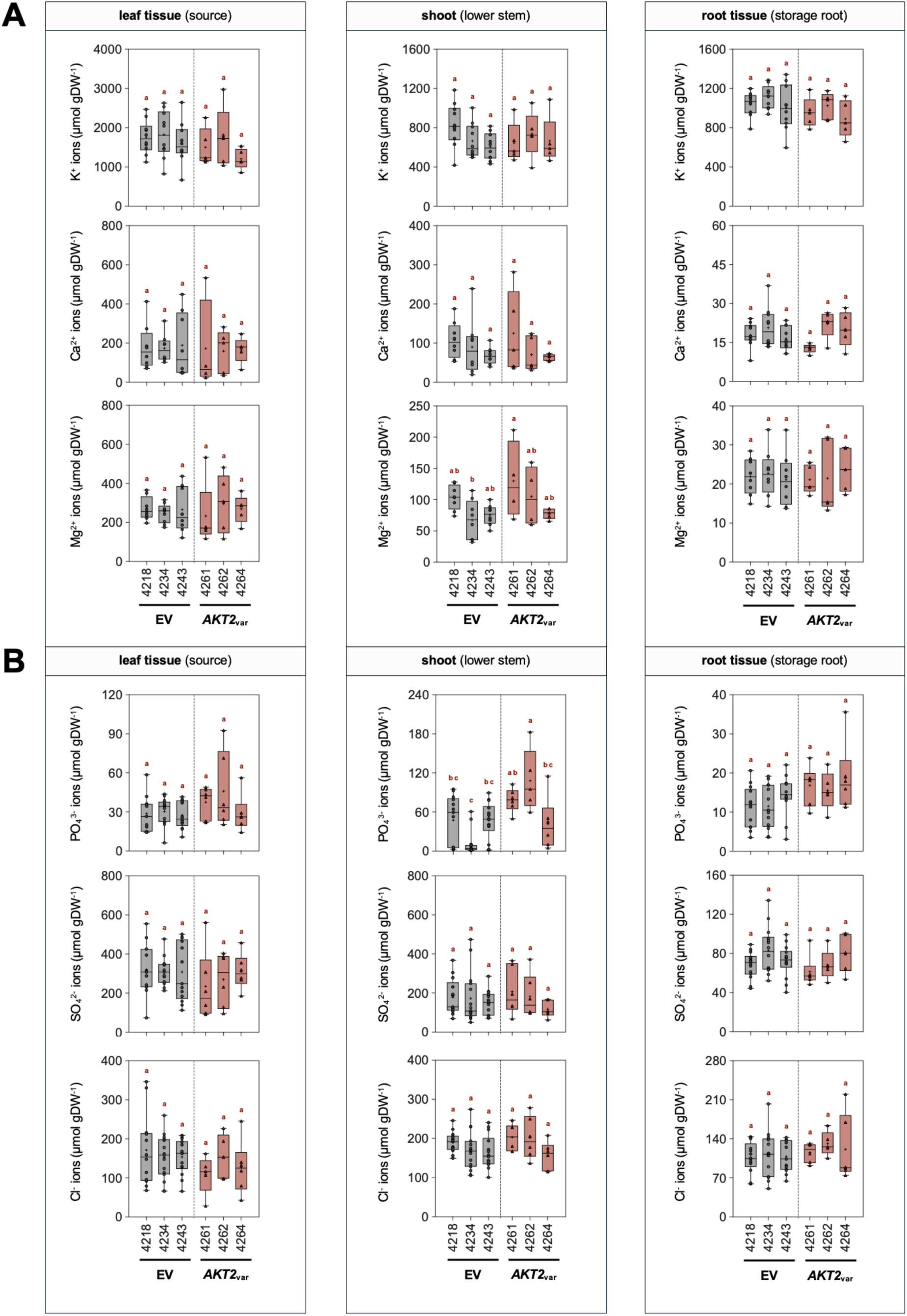
Expression of *AtAKT2_var_* in cassava causes only minor changes in cation and anion distribution in controlled greenhouse experiments. (**A**) Cation contents of potassium (K^+^), calcium (Ca^2+^), and magnesium (Mg^2+^) in leaf tissue, lower stem, and storage root tissue. (**B**) Anion contents as phosphate (PO_4_^3-^), sulphate (SO_4_^2-^), and chloride (Cl^−^) in leaf tissue, lower stem, and storage root tissue. Data in (A) and (B) are from plants of three EV control lines (EV-4218, 4234, and 4243) and three *AKT2_var_* lines (*AKT2_var_*-4261, 4262, and 4264) 19 weeks after planting in soil. The horizontal line in the box plots represents the median and plus (+) the mean. Different lower-case letters indicate statistical significance, as calculated by one-way ANOVA with a post-hoc Tukey HSD test (*p* < 0.05).

**Fig. S4.**
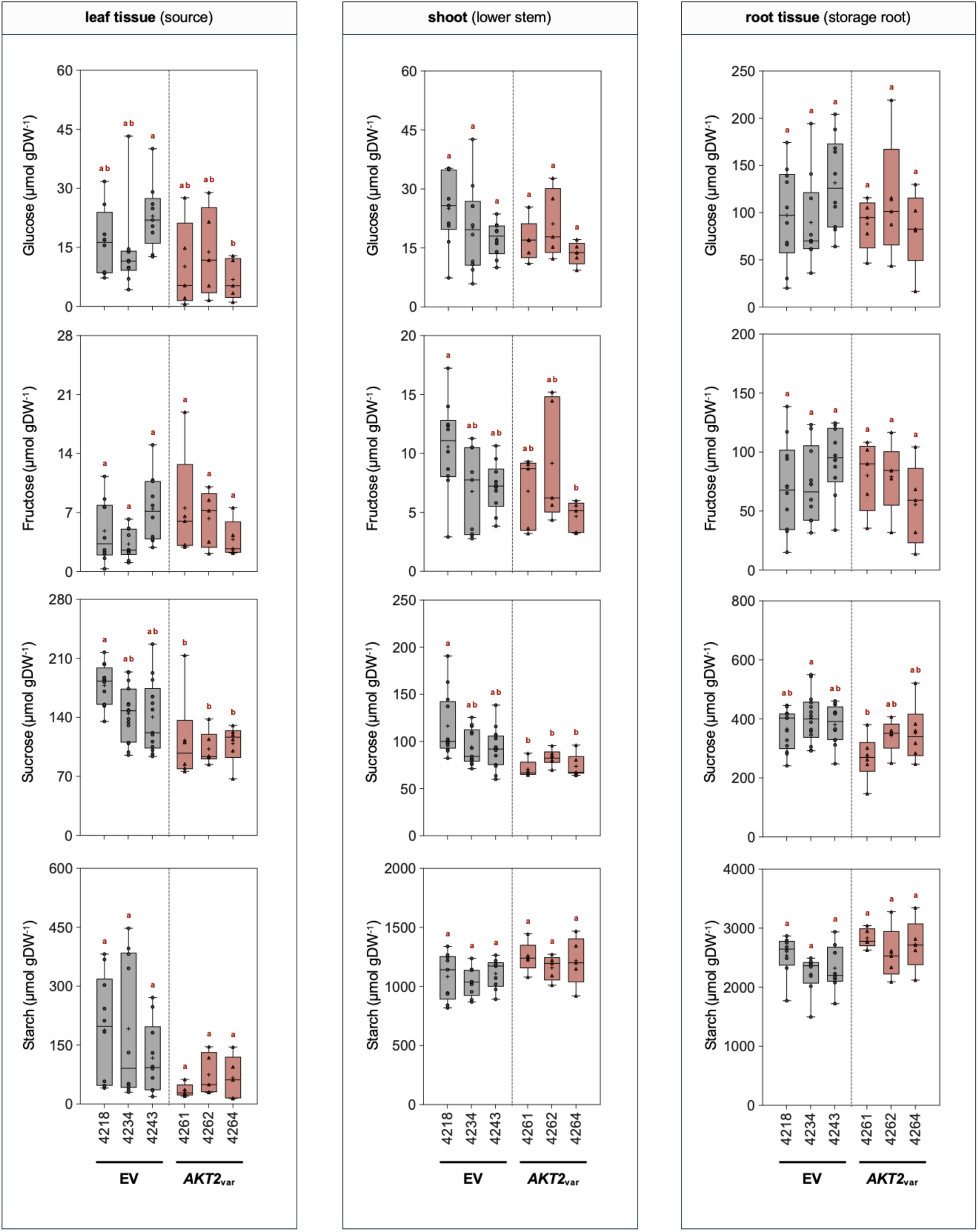
Expression of *AtAKT2_var_* in cassava causes only minor changes in sugar and starch levels in greenhouse experiments. Glucose, fructose, sucrose, and starch concentrations in leaf, lower stem, and storage root tissues of plants from EV control lines (EV-4218, 4234, and 4243), and plants from *AKT2_var_* lines (*AKT2_var_*-4261, 4262, and 4264) 19 weeks after planting in soil. The horizontal line in the box plots represents the median and plus (+) the mean, box borders represent first and third quartiles, whiskers represent the maximum and minimum values, and dots represent single values. Different lower-case letters indicate statistical significance, as calculated by one-way ANOVA with a post-hoc Tukey HSD test (*p* < 0.05).

**Fig. S5.**
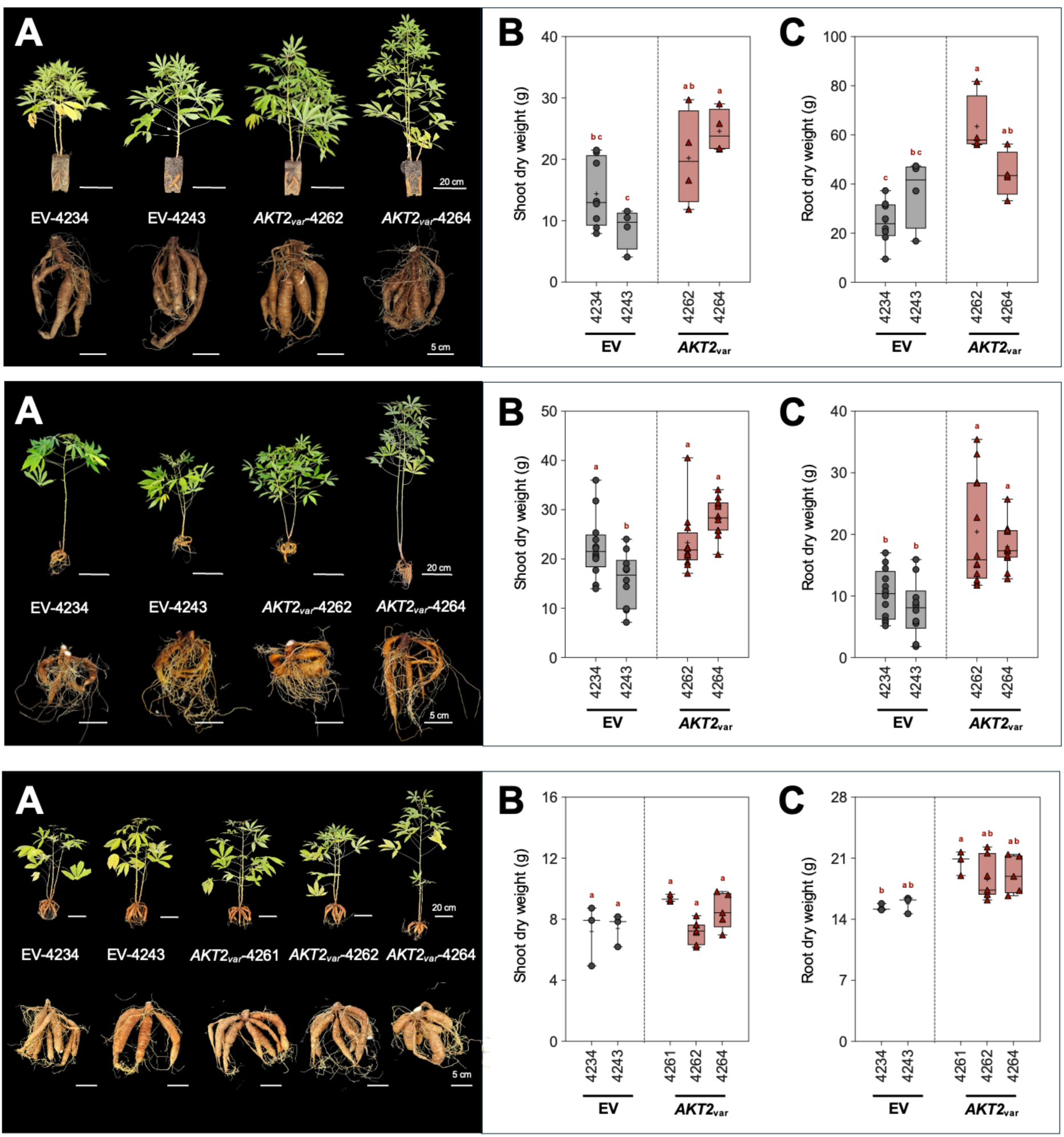
*AtAKT2_var_* expression in cassava enhances shoot and root growth under controlled greenhouse conditions. Data are shown for three replicated experiments. (**A**) Phenotypes of shoots and roots of plants from EV control lines (EV-4234 and 4243), and plants from selected *AKT2_var_* lines (*AKT2_var_*-4261, 4262, and 4264). (**B**) Shoot dry weight and (**C**) root dry weight were measured 19 weeks after planting in soil. The horizontal line in the box plots represents the median and plus (+) the mean, box borders represent first and third quartiles, whiskers represent the maximum and minimum values, and dots represent single values. Different lower-case letters indicate statistical significance, as calculated by one-way ANOVA with a post-hoc Tukey HSD test (*p* < 0.05).

**Fig. S6.**
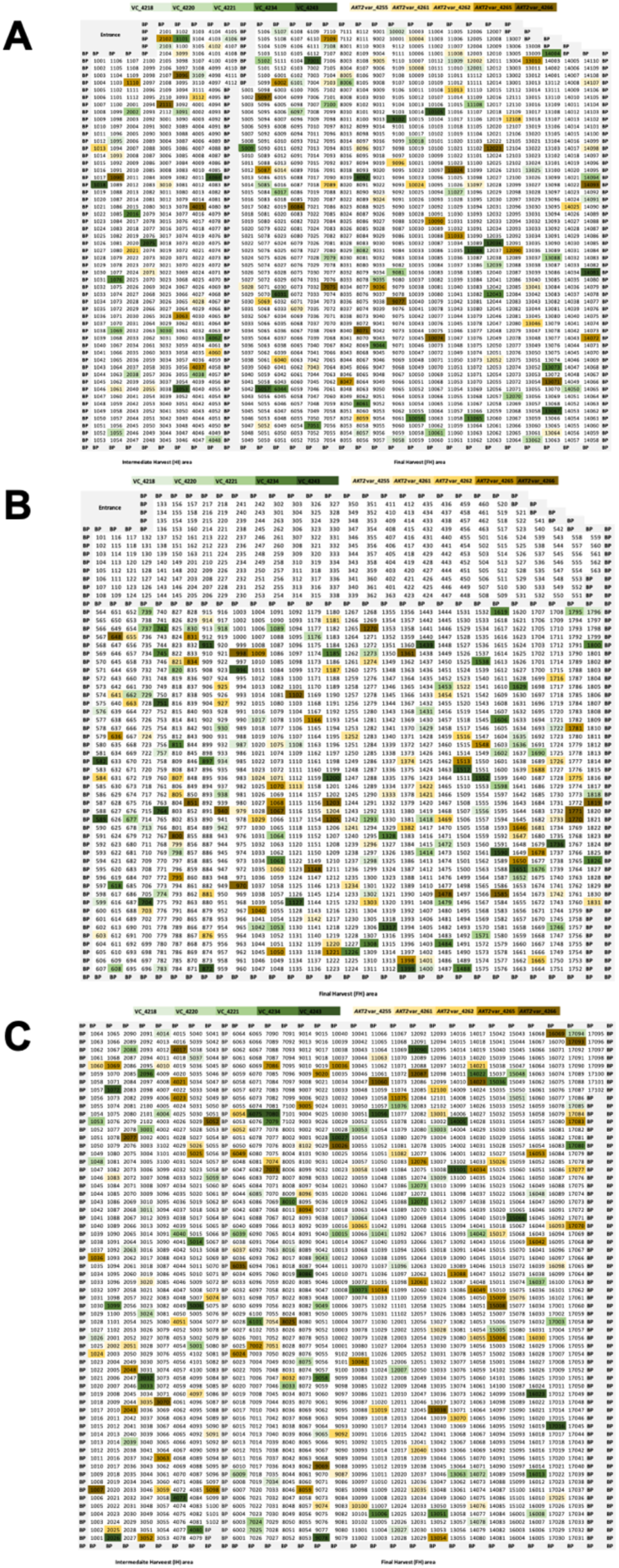
Serpentine randomization of planting for confined field trials in 2022, 2023, and 2024. Serpentine randomization of planting in (**A**) 2022, (**B**) 2023, and (**C**) 2024. Each four-digit number represents a plant grown in that year. Plants of EV lines are shown in shades of green and plants of *AKT2*_var_ lines are shown in shades of orange/brown. Four replications were grown for an intermediate harvest in 2022, 10 replications were grown for the final harvest in 2022. Seventeen replications were grown for the final harvest in 2023. Five replications were grown for an intermediate harvest in 2024, 12 replications were grown for the final harvest in 2024.

**Fig. S7.**
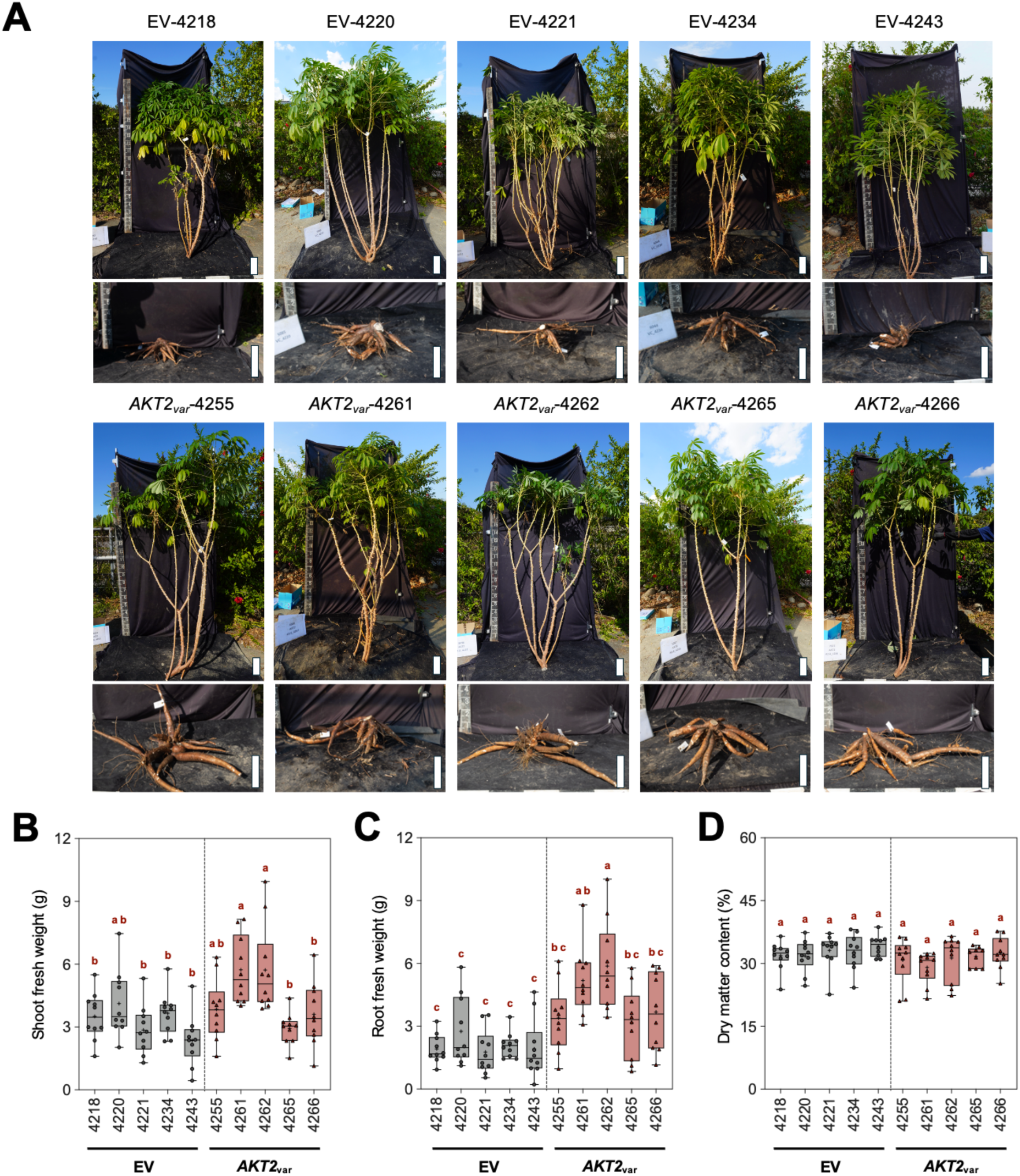
Expression of *AKT2_var_* enhances cassava shoot and root growth under confined field trial conditions in 2022. (**A**) Stems and storage roots of representative plants from EV control and *AKT2_var_*lines. (**B**) Shoot fresh weight, (**C**) root fresh weight, and (**D**) dry matter content was quantified about 9 months (April to December) after planting in 2022. The horizontal line in the box plots represents the median and plus (+) the mean, box borders represent first and third quartiles, whiskers represent the maximum and minimum values, and dots represent single values. Different lower-case letters indicate statistical significance, as calculated by one-way ANOVA with a post-hoc Tukey HSD test (*p* < 0.05). Scale bars are 20 cm.

**Fig. S8.**
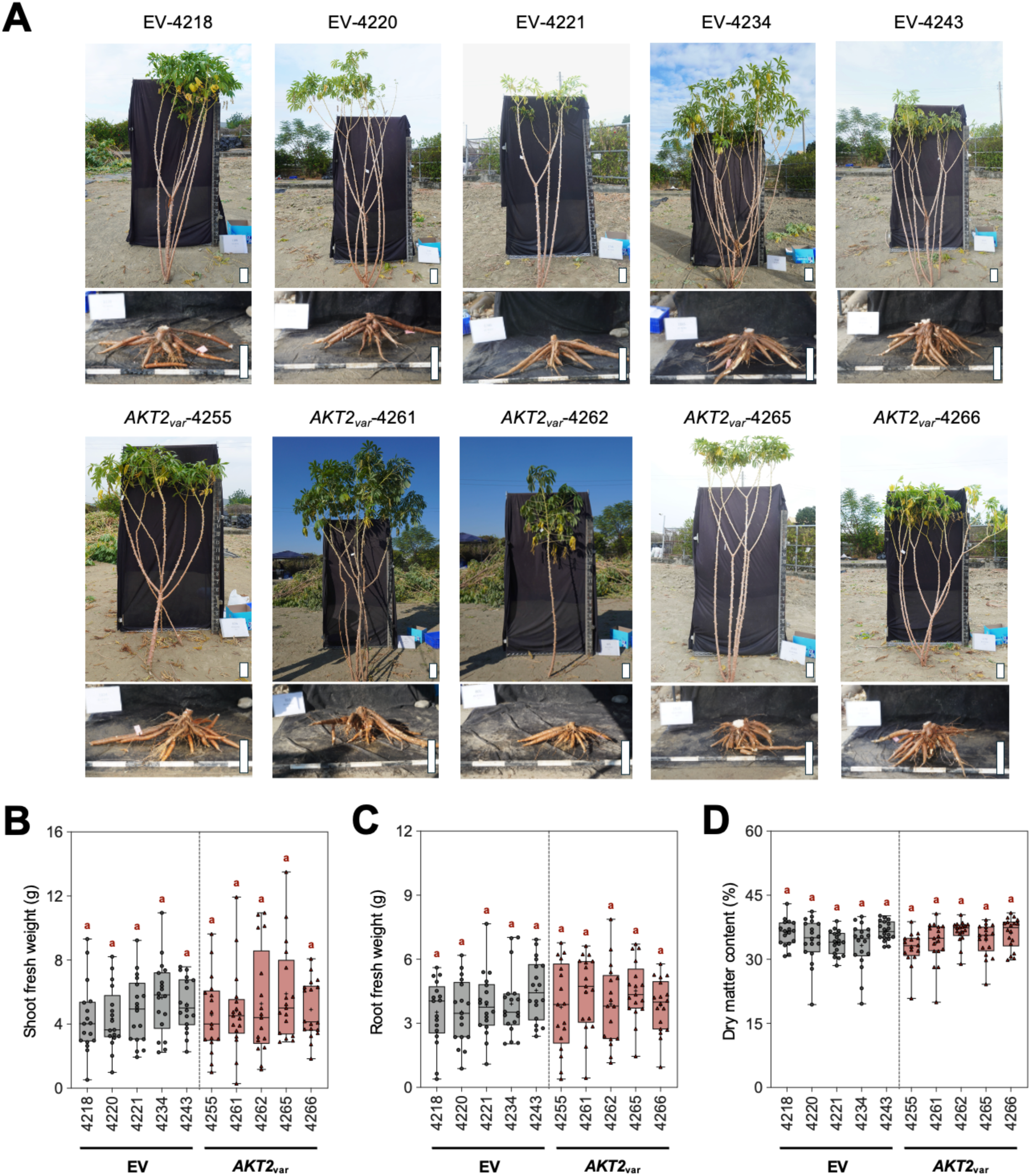
Shoot and root growth of plants from *AKT2_var_* and EV cassava lines under confined field trial conditions in 2023. (**A**) Stems and storage roots of representative plants from EV control and *AKT2_var_*lines. (**B**) Shoot fresh weight, (**C**) root fresh weight, and (**D**) dry matter content was quantified about 9 months (April to December) after planting in 2023. The horizontal line in the box plots represents the median and plus (+) the mean, box borders represent first and third quartiles, whiskers represent the maximum and minimum values, and dots represent single values. Different lower-case letters indicate statistical significance, as calculated by one-way ANOVA with a post-hoc Tukey HSD test (*p* < 0.05). Scale bars are 20 cm.

**Fig. S9.**
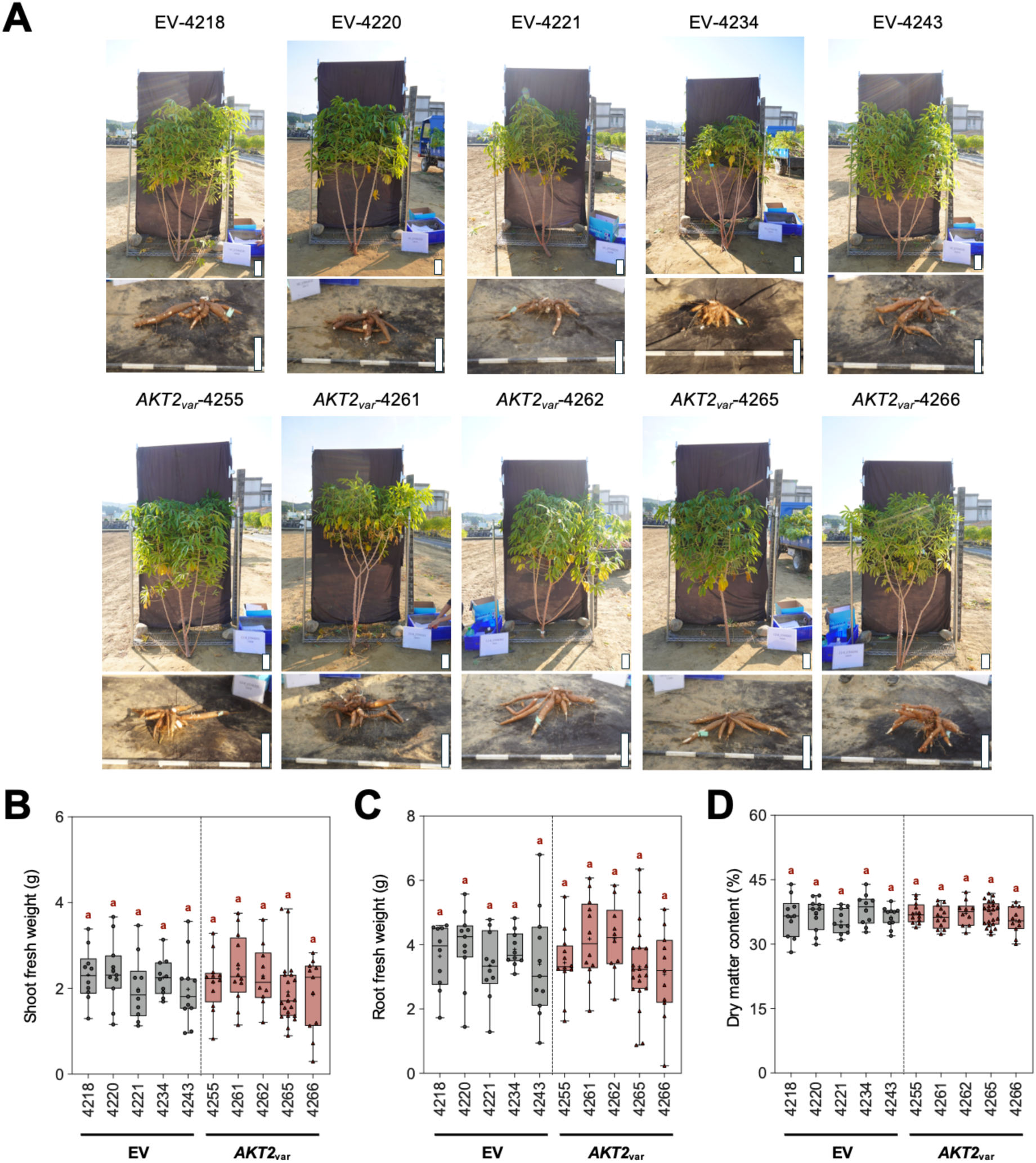
Shoot and root growth of plants from *AKT2_var_* and EV cassava lines under confined field trial conditions in 2024. (**A**) Stems and storage roots of representative plants from EV and *AKT2_var_* lines. (**B**) Shoot fresh weight, (**C**) root fresh weight, and (**D**) dry matter content was quantified about 9 months (April to December) after planting in 2024. The horizontal line in the box plots represents the median and plus (+) the mean, box borders represent first and third quartiles, whiskers represent the maximum and minimum values, and dots represent single values. Different lower-case letters indicate statistical significance, as calculated by one-way ANOVA with a post-hoc Tukey HSD test (*p* < 0.05). Scale bars are 20 cm.

**Fig. S10.**
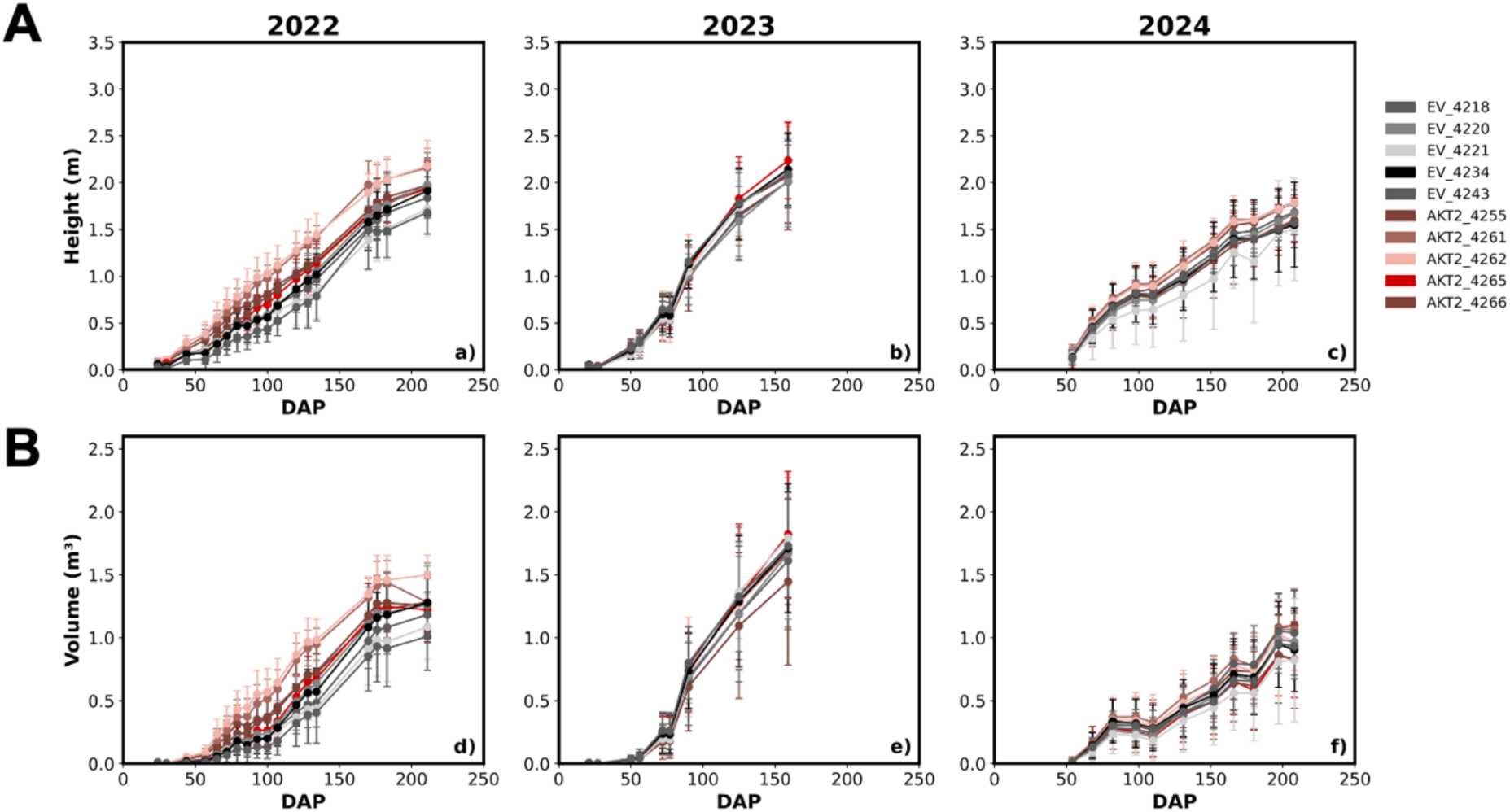
Summary of UAV-phenotyping data recorded during confined field trials. Comparisons between plants from five EV and five *AKT2_var_* lines grown during confined field trials in 2022 to 2024. (**A**) Time course of plant height calculated from UAV data. (**B**) Time course of plant canopy volume calculated from UAV data. UAV = Unmanned aerial vehicle.

**Fig. S11.**
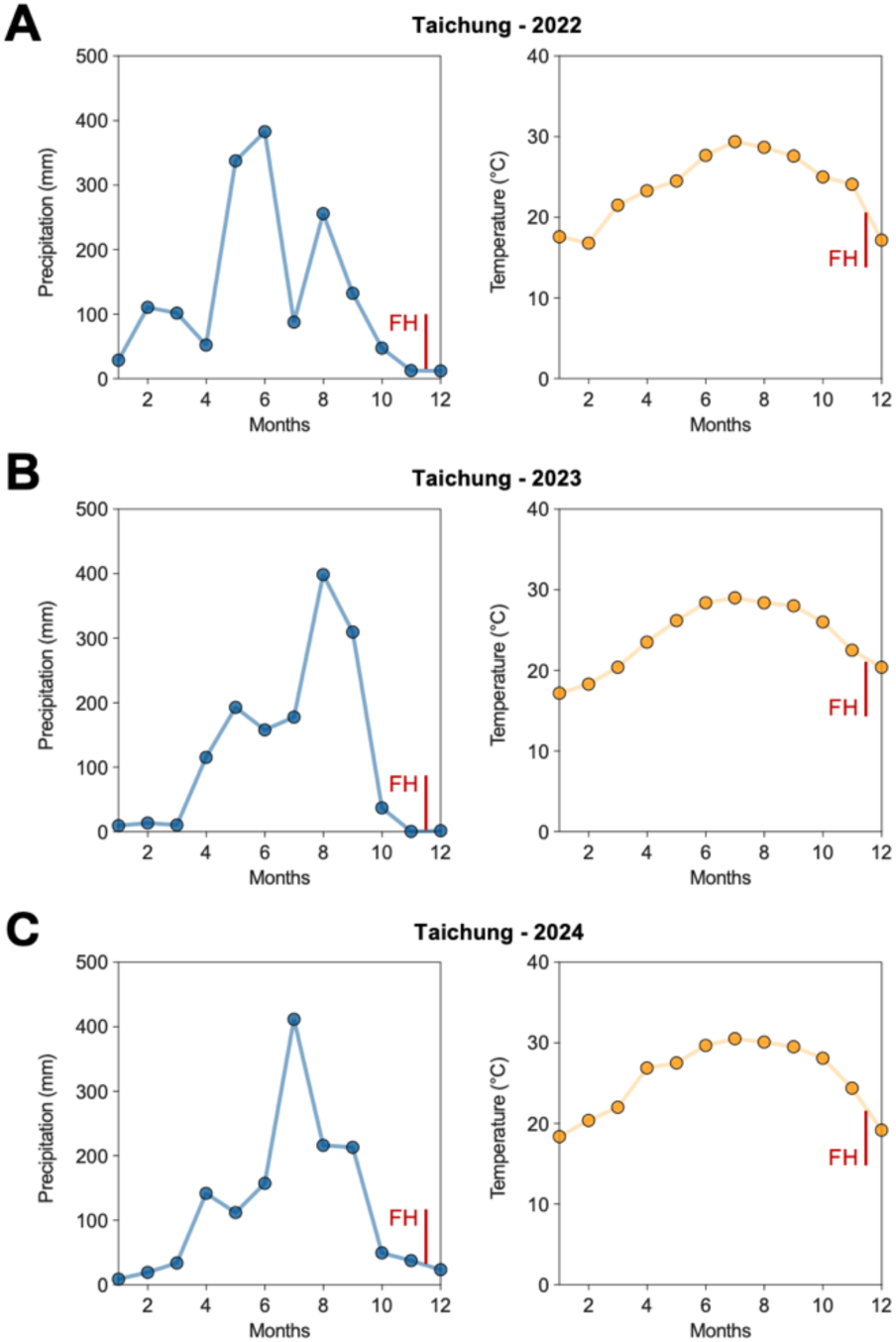
Weather data showing differences in rainfall during the 2022, 2023, and 2024 confined field trials. Precipitation data and the temperature measurements from field trials in (**A**) 2022, (**B**) 2023, and (**C**) 2024. Harvests were completed about nine months after transfer of cassava plantlets to the field.

**Fig. S12.**
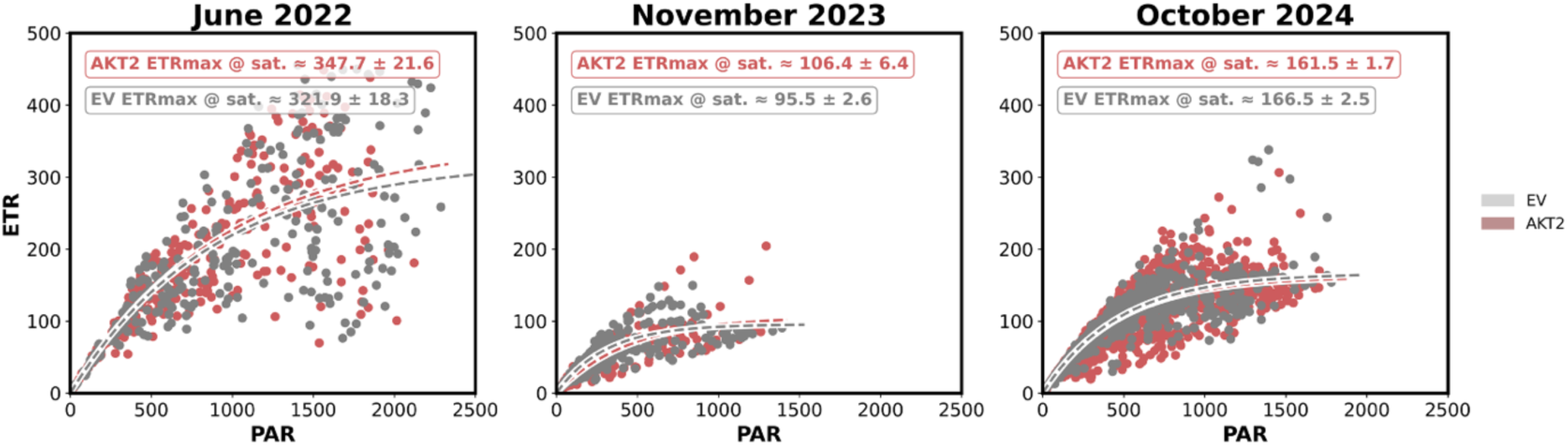
Summary of electron transport rate (ETR) data obtained during confined field trials. Recorded ETR values at different Photosynthetically Active Radiation (PAR) levels are shown for plants from EV and *AKT2_var_* lines grown during confined field trials in 2022 to 2024. The dashed lines represent the integrated curves calculated from all values obtained from EV and *AKT2_var_* plants. Data from 2022 was collected in June from approximately 3-month-old cassava plants, data from 2023 was collected in November from approximately 8-month-old cassava plants, and data from 2024 was collected in October from approximately 7-month-old cassava plants.

**Fig. S13.**
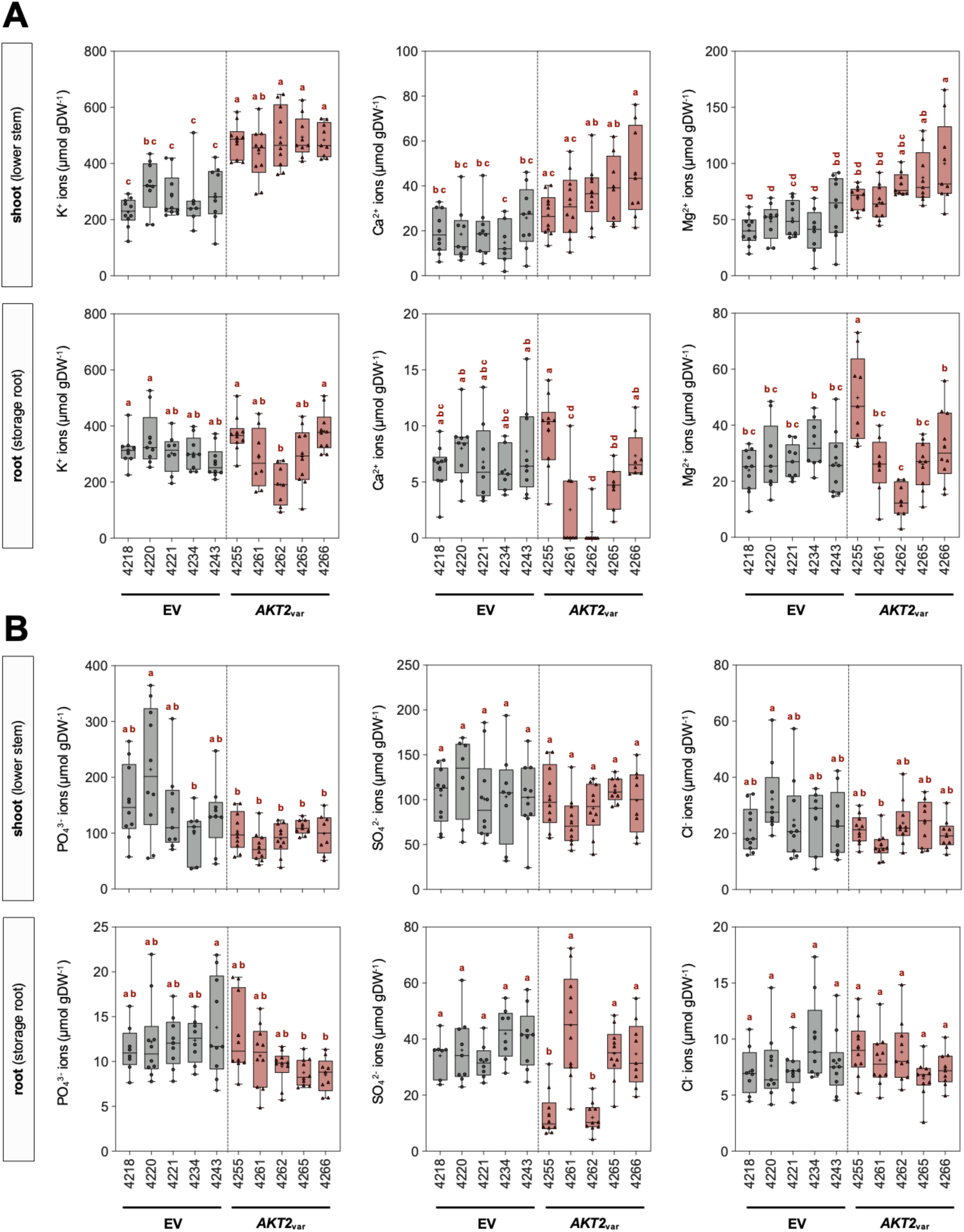
*AKT2_var_* expression in cassava causes alterations in cation and anion concentrations in confined field trials in 2022. (**A**) Cation contents of potassium (K^+^), calcium (Ca^2+^), and magnesium (Mg^2+^) in lower stem and storage root tissues. (**B**) Anion contents of phosphate (PO_4_^3-^), sulphate (SO_4_^2-^), and chloride (Cl^−^) in lower stem and storage root tissues. Data are shown for plants from EV lines (EV-4218, 4234, and 4243) and *AKT2_var_* lines (*AKT2_var_*-4261, 4262, and 4264) that were harvested about nine months (April to December) after planting. The horizontal line in the box plots represents the median and plus (+) the mean. Different lower-case letters indicate statistical significance, as calculated by one-way ANOVA with a post-hoc Tukey HSD test (*p* < 0.05).

**Fig. S14.**
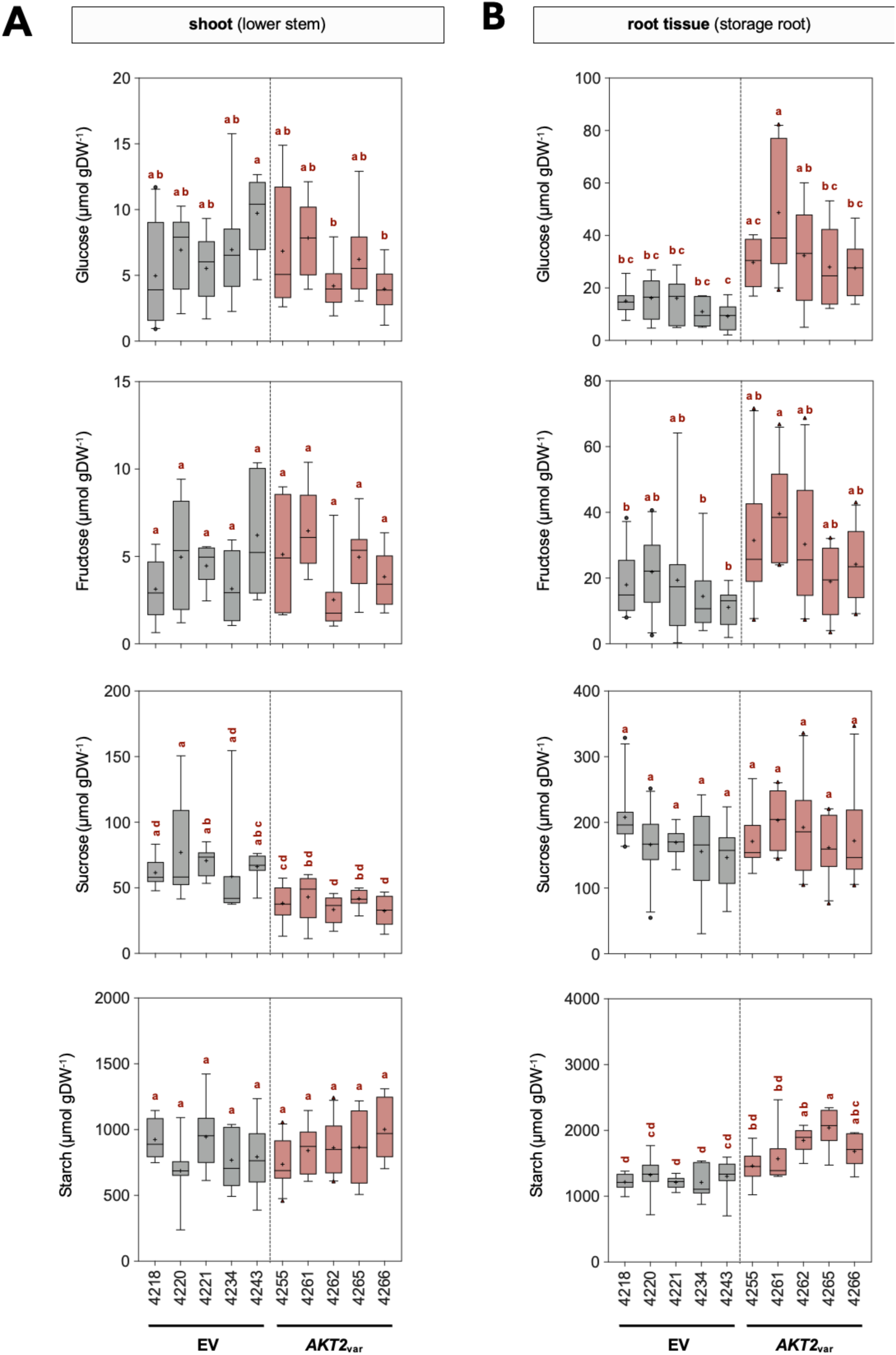
Expression of *AKT2_var_* in cassava leads to changes in sugar concentrations in 2022 confined field trials. Glucose, fructose, sucrose, and starch concentrations in (**A**) shoot and (**B**) storage root tissues. Data are shown for plants from EV lines (EV-4218, 4234, and 4243) and *AKT2_var_* lines (*AKT2_var_*-4261, 4262, and 4264) about nine months (April to December) after planting. The horizontal line in the box plots represents the median and plus (+) the mean. Different lower-case letters indicate statistical significance, as calculated by one-way ANOVA with a post-hoc Tukey HSD test (*p* < 0.05).

**Fig. S15.**
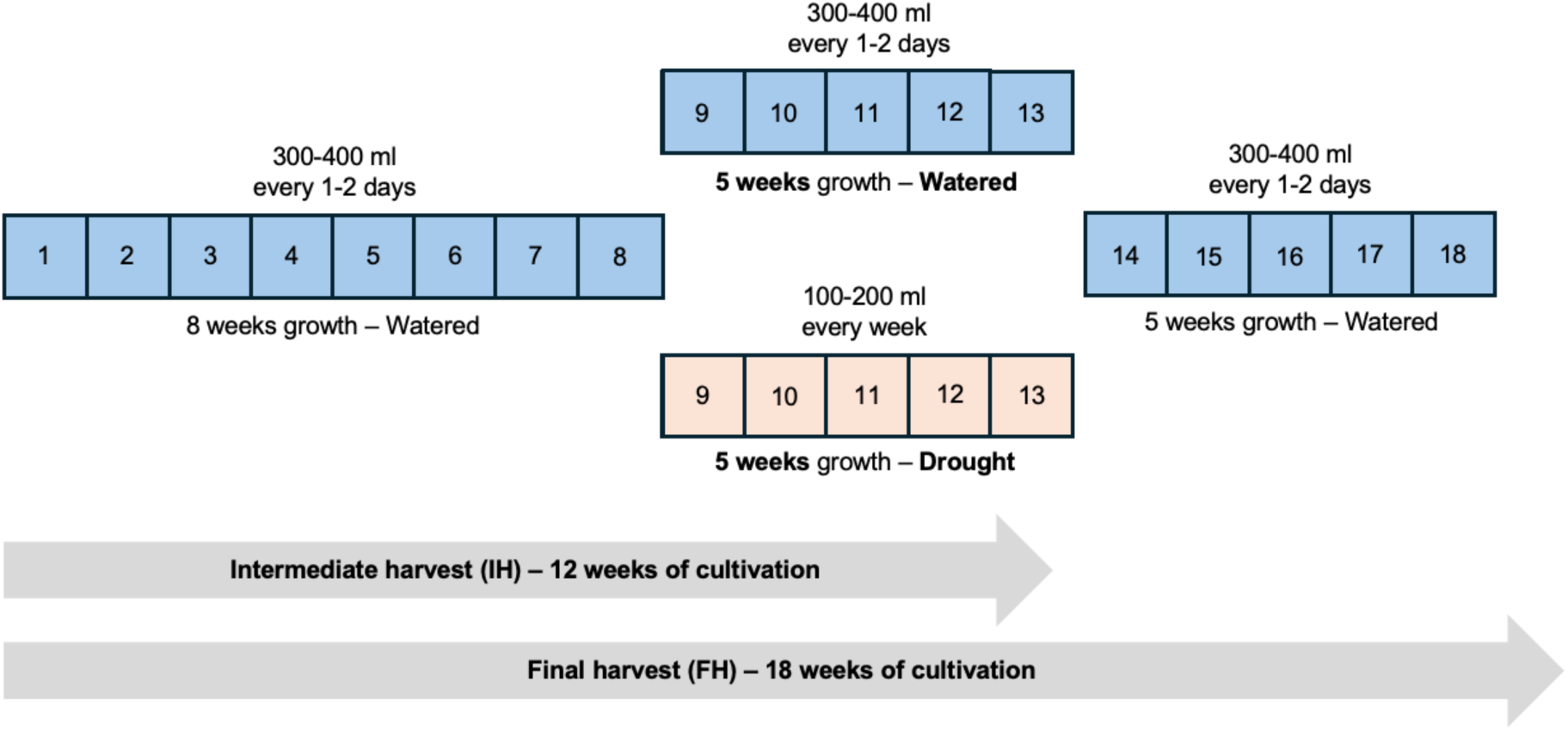
Schematic representation of the cultivation process to induce drought stress in cassava under greenhouse conditions. To induce drought stress in the EV and *AKT2_var_* lines, all plants were first grown under controlled watered conditions for 8 weeks. This was followed by a five-week drought stress period during which half of the plants were watered only once a week, while control plants were continued to be watered daily. An intermediate harvest was carried out after five weeks of drought, after which all plants were again watered daily. The final harvest was carried out after a further five weeks of growth.

**Fig. S16.**
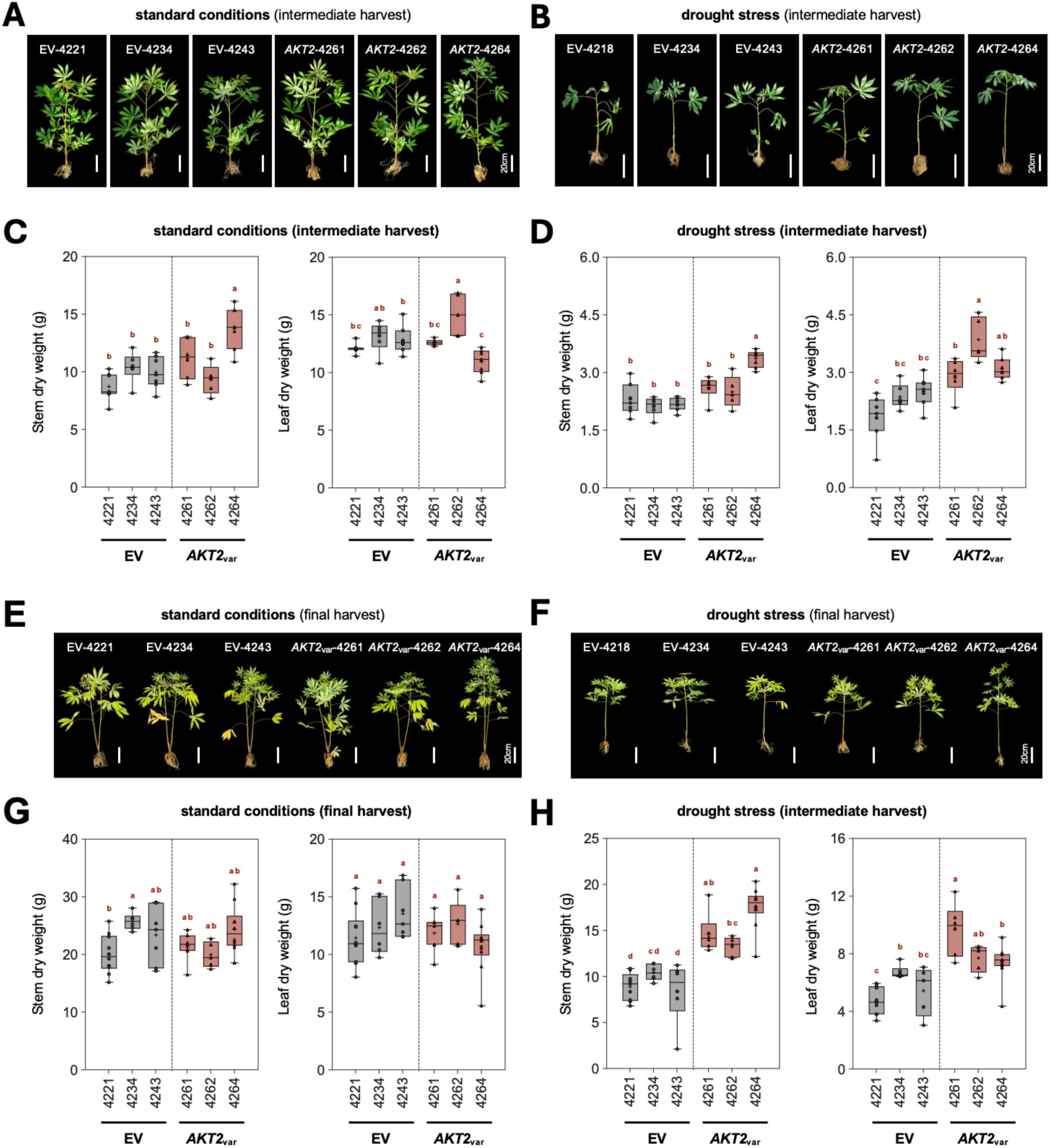
*AKT2_var_* expression not only enhances cassava growth under standard conditions, but also significantly improves the response to drought stress. Plants from EV and *AKT2_var_* lines were grown, treated for drought and sampled according to the schedule shown in Fig. S15. (**A, B**) Phenotypes of shoots and roots from plants of EV lines (EV-4221, 4234, and 4243) and *AKT2_var_* lines (*AKT2_var_*-4261, 4262, and 4264) at intermediate harvest (IH). (**C, D**) Stem, leaf, and root dry weights were measured at intermediate harvest under either standard or drought stress conditions. (**E, F**) Phenotypes of shoots and roots from plants of EV lines (EV-4221, 4234, and 4243), and *AKT2_var_* lines (*AKT2_var_*-4261, 4262, and 4264) at final harvest (FH). (**G, H**) Stem, leaf, and root dry weights were measured at final harvest under either control, or after drought stress conditions. The horizontal line in the box plots represents the median and plus (+) the mean. Different lower-case letters indicate statistical significance, as calculated by one-way ANOVA with a post-hoc Tukey HSD test (*p* < 0.05).

**Fig. S17.**
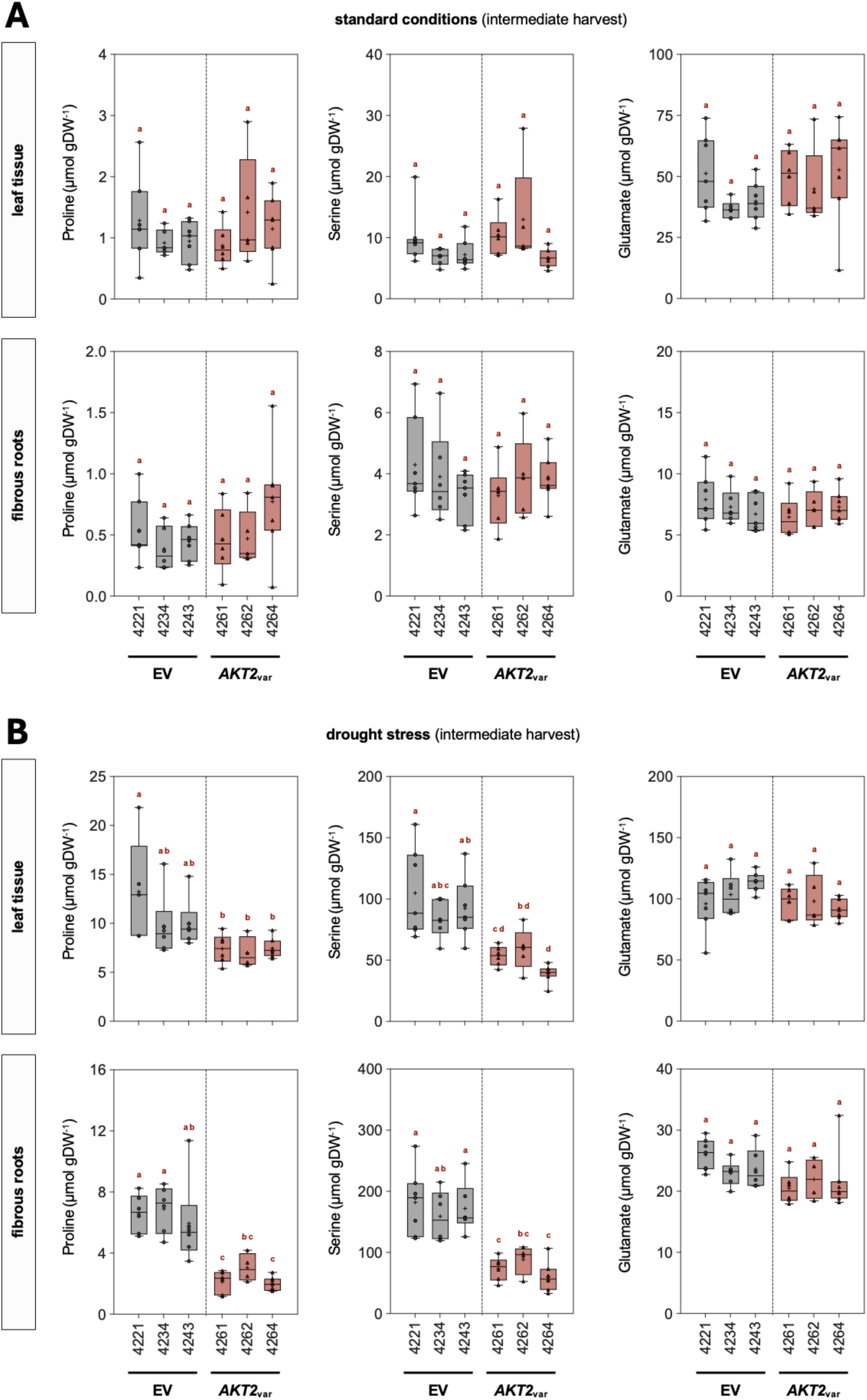
*AKT2_var_* lines have reduced levels of stress-related amino acids during drought stress. Plants from EV and *AKT2_var_* lines were grown, treated for drought and sampled according to the schedule shown in Fig. S15. The amino acids proline, serine, and glutamate were measured in leaf tissue and fibrous roots of plants from all analysed EV lines (EV-4221, 4234, and 4243) and *AKT2_var_* lines (*AKT2_var_*-4261, 4262, and 4264) under (**A**) control conditions (**B**) or during the drought stress phase at intermediate harvest. The horizontal line in the box plots represents the median and plus (+) the mean. Different lower-case letters indicate statistical significance, as calculated by one-way ANOVA with a post-hoc Tukey HSD test (*p* < 0.05).

**Fig. S18.**
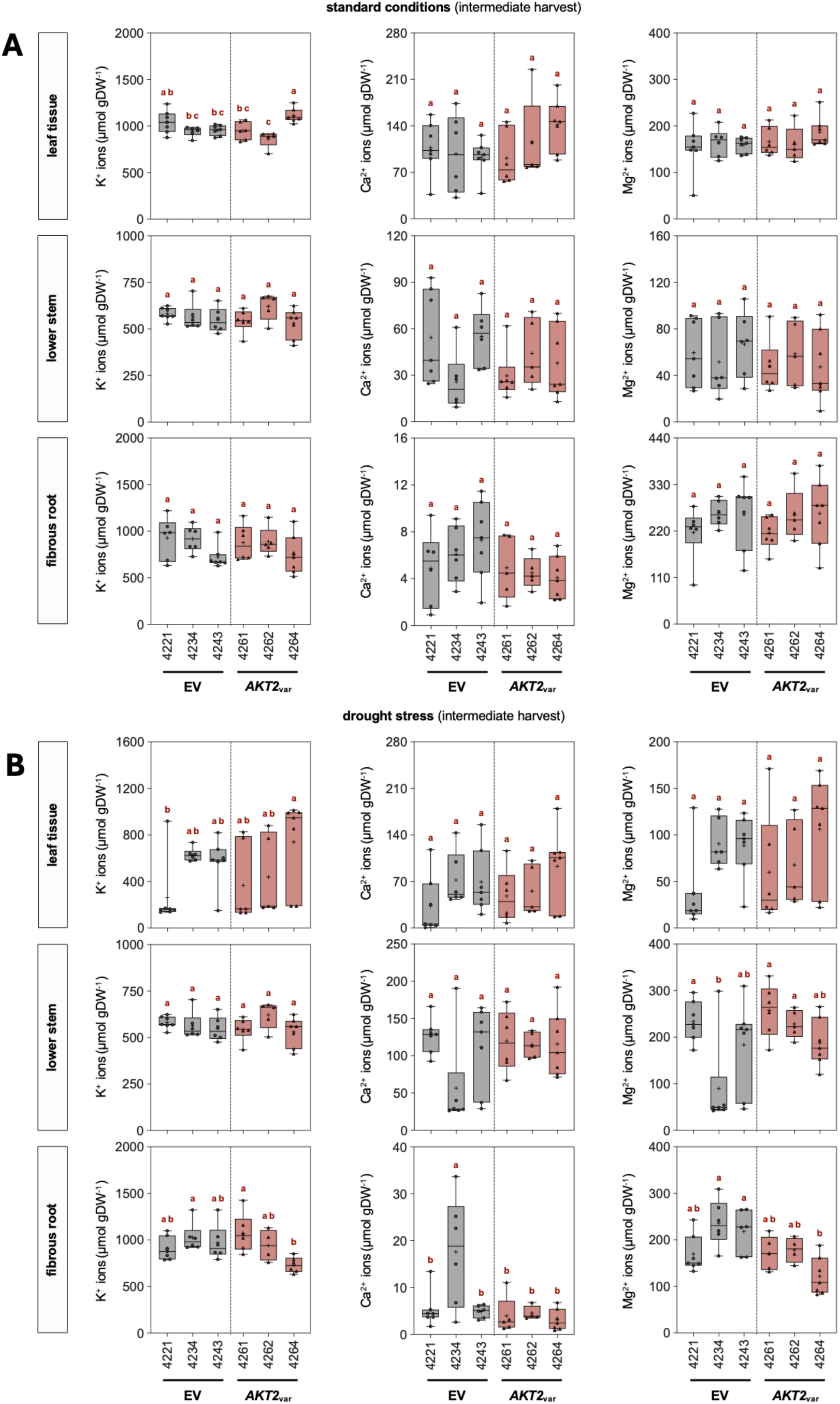
*AKT2_var_* expression in cassava does not cause changes in cation distribution during periodic drought stress. Concentrations of potassium (K^+^), calcium (Ca^2+^), and magnesium (Mg^2+^) in leaf, lower stem, and fibrous root tissues of plants from EV lines (EV-4221, 4234, and 4243) and *AKT2_var_* lines (*AKT2_var_*-4261, 4262, and 4264) under (**A**) control conditions or (**B**) during periodic drought stress. The horizontal line in the box plots represents the median and plus (+) the mean. Different lower-case letters indicate statistical significance, as calculated by one-way ANOVA with a post-hoc Tukey HSD test (*p* < 0.05).

**Fig. S19.**
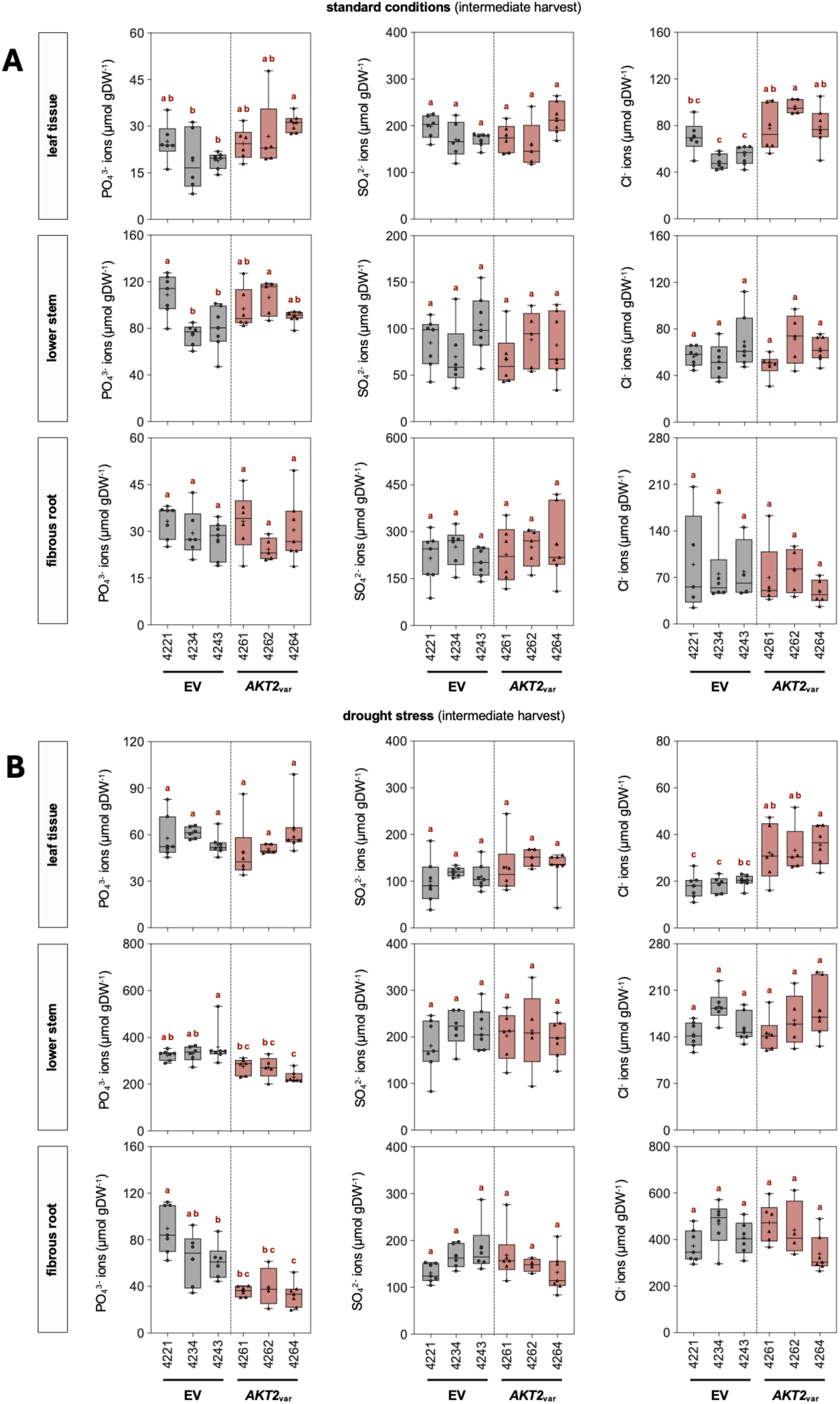
*AKT2_var_* expression in cassava causes only minor changes in anion distributions during periodic drought stress. Shown are cation contents of phosphate (PO_4_^3-^), sulphate (SO_4_^2-^), and chloride (Cl^−^) in leaf, lower stem, and fibrous root tissues of plants from EV lines (EV-4221, 4234, and 4243) and *AKT2_var_* lines (*AKT2_var_*-4261, 4262, and 4264) under (**A**) control conditions or (**B**) during periodic drought stress. The horizontal line in the box plots represents the median and plus (+) the mean. Different lower-case letters indicate statistical significance, as calculated by one-way ANOVA with a post-hoc Tukey HSD test (*p* < 0.05).

**Fig. S20.**
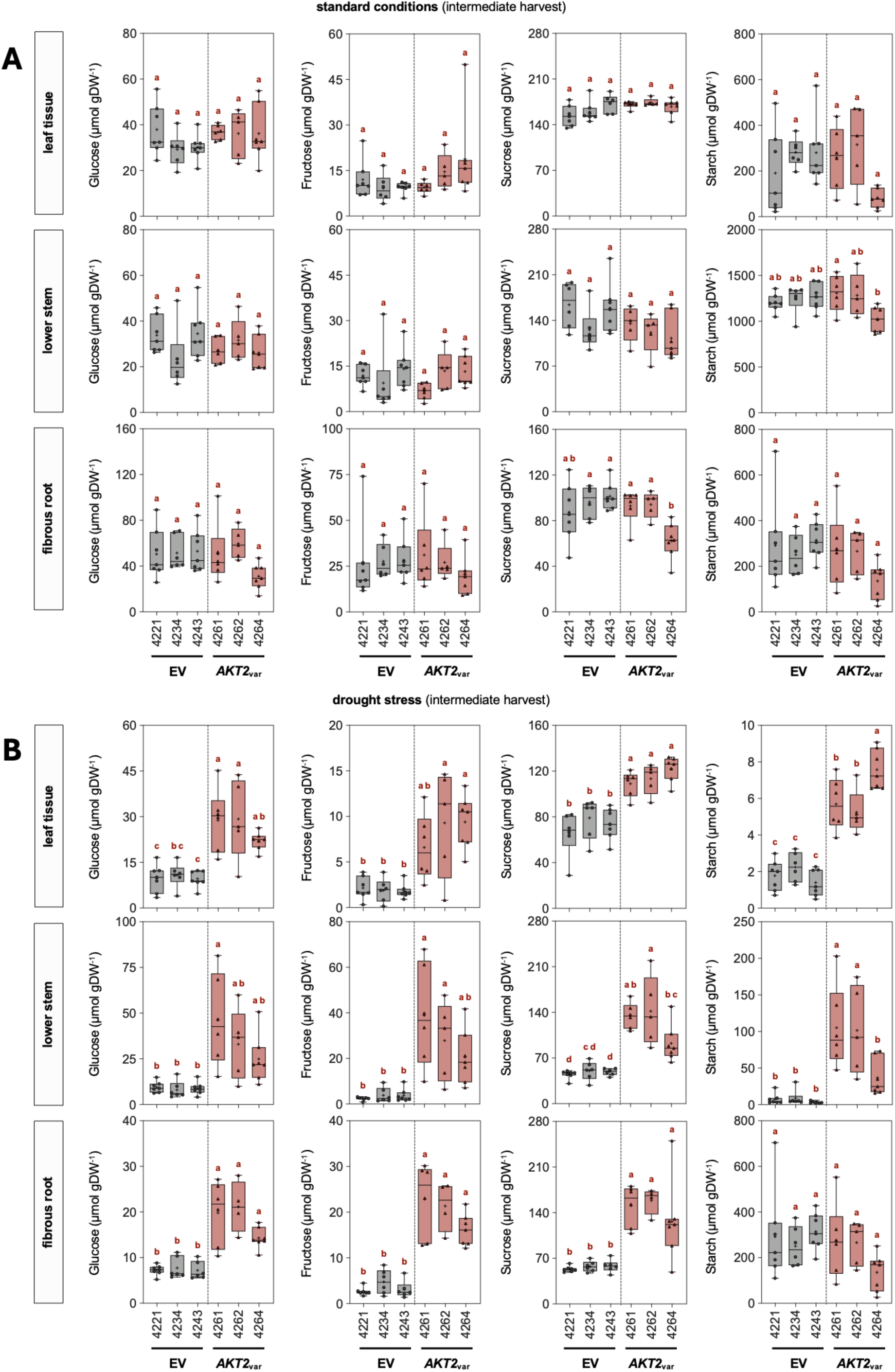
*AKT2_var_* expression in cassava leads to increased carbohydrate levels during periodic drought stress. Glucose, fructose, sucrose, and starch concentrations in leaf, lower stem, and fibrous root tissues of plants from EV lines (EV-4221, 4234, and 4243) and *AKT2_var_* lines (*AKT2_var_*-4261, 4262, and 4264) under (**A**) control conditions or (**B**) during periodic drought stress. The horizontal line in the box plots represents the median and plus (+) the mean. Different lower-case letters indicate statistical significance, as calculated by one-way ANOVA with a post-hoc Tukey HSD test (*p* < 0.05).

**Fig. S21.**
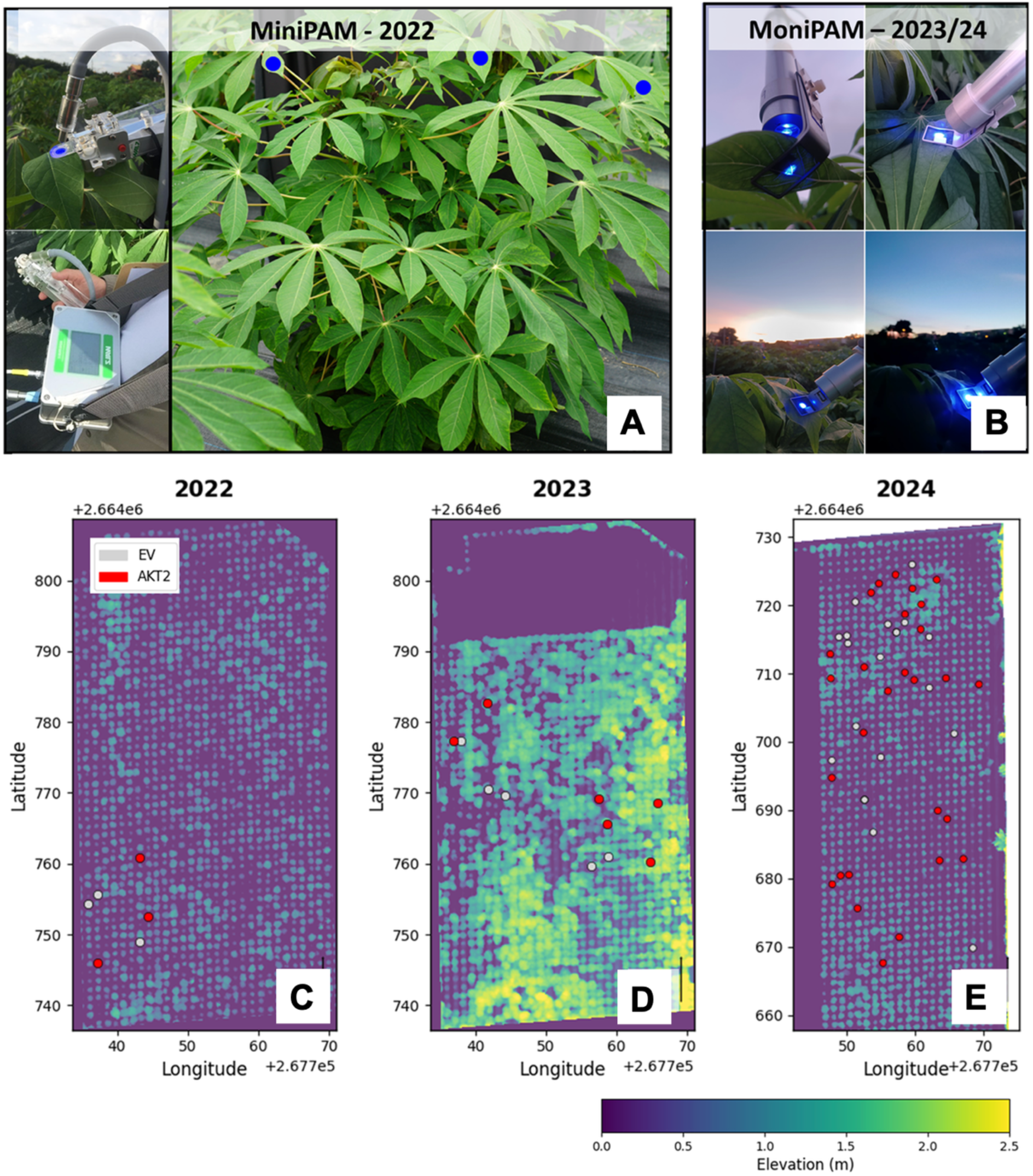
Photosynthetic phenotyping and elevation mapping of cassava genotypes across three seasons. (**A**) Mini-PAM fluorometry setup used in the 2022 field campaign. (**B**) Moni-PAM system applied in the 2023/24 season for high-throughput diurnal measurements. (**C-E**) UAV-derived elevation maps of selected field blocks acquired on July 7, 2022 (2022), July 25, 2023 (2023), and August 28, 2024 (2024). The colour scale represents relative surface elevation (0.0–2.5 m), derived from canopy surface models. Plants used for photosynthesis are indicated by red markers belonging to the *AKT2_var_* genotype group, while grey markers indicate EV (empty vector) control lines.

### Supplementary Tables

**Table S1:**
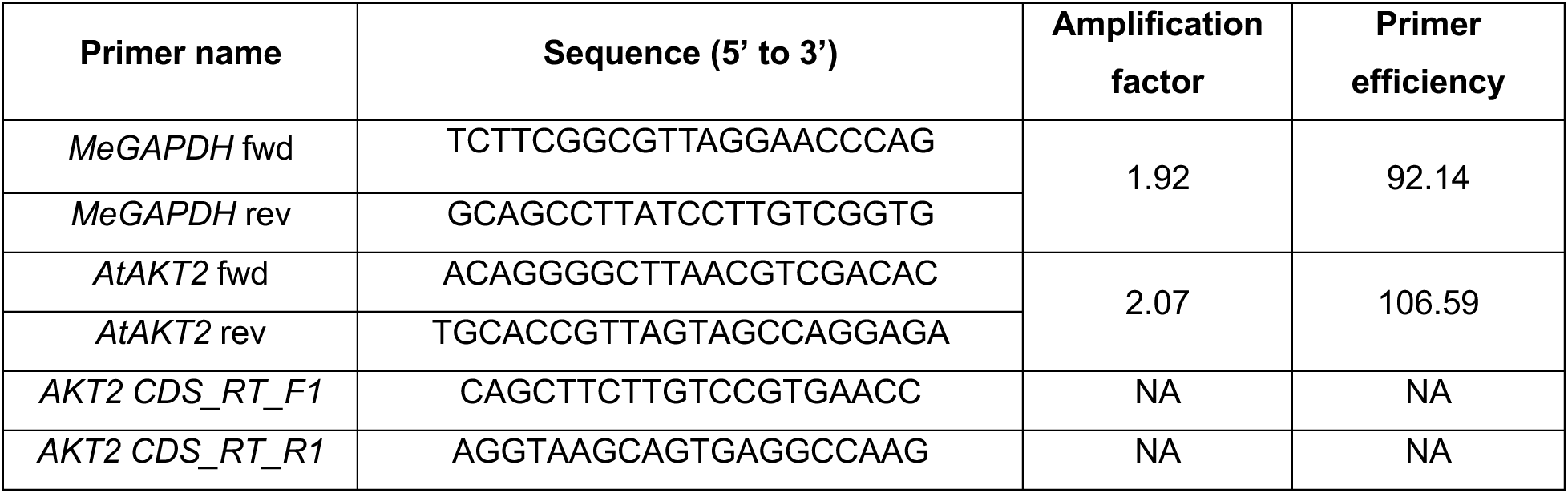
Primer list for quantitative or semiquantitative reverse transcription PCR.

### Supplemental Data

**Supplementary Data 1: Summary of data used in this study**

**Supplementary Data 2: Plasmids used in this study**

